# Regulation of Leucine-Rich Repeat Kinase 2 by inflammation and IL-4

**DOI:** 10.1101/2024.04.29.591170

**Authors:** Dina Dikovskaya, Rebecca Pemberton, Matthew Taylor, Anna Tasegian, Karolina Zenevicuite, Esther M. Sammler, Andrew J.M. Howden, Dario R. Alessi, Mahima Swamy

## Abstract

Mutations in Leucine-Rich Repeat protein Kinase 2 (LRRK2) are associated with Parkinson’s Disease (PD) and Crohn’s Disease (CD), but the regulation of LRRK2 during inflammation remains relatively unexplored. Here we developed a flow cytometry-based assay to assess LRRK2 activity in individual cells and created an EGFP-Lrrk2-knock-in reporter mouse to analyse cell-specific LRRK2 expression. Using these tools, we catalogued LRRK2 levels and activity in splenic and intestinal immune cells. Inflammation increased LRRK2 expression and activity in B-cells, immature neutrophils and immature monocytes, but decreased these in dendritic cells and eosinophils. In mature neutrophils, inflammation stimulated activity but reduced LRRK2 expression. A kinase-activating PD-associated LRRK2-R1441C mutation exacerbated inflammation-induced activation of LRRK2 specifically in monocytes and macrophages without affecting LRRK2 levels. Finally, we identified IL-4 as a novel factor that upregulated LRRK2 expression and activity in B-cells, replicating inflammatory effects observed *in vivo*. Our findings provide valuable new insights into the regulation of the LRRK2 pathway in immune cells, crucial for understanding LRRK2 and its therapeutic potential in inflammatory diseases such as CD.

## Introduction

Leucine-Rich Repeat protein Kinase 2 (LRRK2) is a large multidomain protein that combines kinase and GTPase enzymatic activities, involved in regulation of multiple intracellular processes such as lysosomal maintenance and functioning, vesicular trafficking (Kuwahara & Iwatsubo, 2020; Roosen & Cookson, 2016), mitophagy (Singh *et al*, 2021), inflammasome activation (Liu *et al*, 2017a) as well as several other innate immune pathways (Ahmadi Rastegar & Dzamko, 2020). Many LRRK2 functions are thought to be mediated by its kinase substrates Rab GTPases (Rab1, Rab3, Rab5, Rab8, Rab10, Rab12, Rab29, Rab35 and Rab43) (Alessi & Pfeffer, 2024). Mutations in LRRK2 are causal for Parkinson’s disease (PD) (Paisan-Ruiz *et al*, 2004; Zimprich *et al*, 2004) and genetically associated with several inflammatory diseases including Crohn’s Disease (CD) (Hui *et al*, 2018; Witoelar *et al*, 2017), leprosy (Fava *et al*, 2016; Wang *et al*, 2015) and systemic lupus erythematosus (Zhang *et al*, 2017). Most of the well characterised disease-associated mutations in LRRK2 cluster within either its kinase domain or a ROC-COR domain and stimulate kinase activity (Kalogeropulou *et al*, 2022), making LRRK2 kinase an attractive drug target. In animal testing studies including non-human primates, high doses of LRRK2 kinase inhibitors that ablate pathway activity revealed lung and kidney abnormalities that are likely due to on-target effects as these are recapitulated in the LRRK2 knock-out phenotype (Araki *et al*, 2018; Wojewska & Kortholt, 2021). However, a recent Phase-1 clinical study with an inhibitor (DNL201) that suppressed LRRK2 pathway activity by ∼70%, concluded that this compound was safe and well tolerated, warranting further clinical development of LRRK2 inhibitors as a therapeutic modality for PD (Jennings *et al*, 2022).

However, the potential effect of LRRK2 inhibition on specific cell types and their functions in the context of inflammatory disease is not well understood. LRRK2 is highly expressed in select human immune cells, including neutrophils, monocytes and B-cells, and its level has been reported to be further increased in B-cells, T-cells and monocytes after prolonged IFNγ stimulation (Ahmadi Rastegar *et al*, 2022; Cook *et al*, 2017; Gardet *et al*, 2010; Hakimi *et al*, 2011; Kuss *et al*, 2014; Thevenet *et al*, 2011). While the role of LRRK2 and the consequences of LRRK2 inhibition have been extensively studied in monocytes and macrophages (Dzamko *et al*, 2015; Herbst *et al*, 2020; Lee *et al*, 2020; Thevenet *et al*., 2011; Yadavalli & Ferguson, 2023), little is known about other cells with active LRRK2. Furthermore, the expression and functions of LRRK2 in intestinal immune cells, relevant to CD, are unknown. This knowledge is critical to understand how LRRK2 inhibitors impact immune cells and their ability to trigger and mediate inflammatory responses.

Here, to establish the landscape of LRRK2 activity among different immune cell types relevant for CD, we developed and validated a flow cytometry-based assay that measures LRRK2-dependent phosphorylation of its substrate Rab10 with single cell resolution. We combined it with a newly developed EGFP-Lrrk2-reporter mouse to better quantify the relative expression levels of LRRK2 protein in mouse splenic and intestinal immune cells and determine how these are affected by intestinal tissue environment and inflammation. We also demonstrate that the PD-associated R1441C mutation in LRRK2 not only enhances the basal LRRK2 activity in most tested LRRK2-expressing immune cells but also exacerbates eosinophils, immature monocytes and macrophages response to inflammation. We further demonstrate that in B-cells, inflammation-driven increases in level and activity of LRRK2 can be recapitulated *in vitro*, accompanied by a 3-fold increase in Lrrk2 mRNA. We identify Interleukin-4 (IL-4) as the cytokine responsible for inducing and activating LRRK2 in B-cells.

## Results

### 1. Development of a flow-cytometric assay for LRRK2 activity in tissues

To identify cells that express LRRK2 in intestinal tissues, the primary site affected by CD, we first attempted to use immunofluorescence (IF) in frozen or paraffin-embedded intestinal preparations from wild-type (WT) mice, controlled by analogous staining in mice deficient for LRRK2 (LRRK2-KO) to establish the specificity of staining. Despite previous reports of successful detection of LRRK2 in mouse brain and gut tissues by IF and IHC (Davies *et al*, 2013; West *et al*, 2014; Zhang *et al*, 2015), we were unable to detect LRRK2-specific staining in mouse tissues with several published anti-LRRK2 antibodies, including c41-2, the most commonly used antibody for LRRK2 in IF. Indeed, the c41-2-elicited signal within ileal lamina propria (**Suppl. Fig. 1A,** left panels), in Peyer’s patches rich in immune cells (**Suppl. Fig. 1A**, second panel) or in lung (**Suppl. Fig. 1B**) was not diminished in tissues obtained from LRRK2-KO mice. To assess the specificity of c41-2 for LRRK2 in IF in a murine cell line, we analysed a mouse small intestinal cell line, MODE-K, that endogenously express high levels of LRRK2, and a LRRK2-deficient MODE-K clone made by CRISPR deletion. Immunoblotting with the highly specific LRRK2 N241A/34 antibody revealed loss of LRRK2 and loss of phosphorylation of LRRK2 kinase substrate Rab10 at residue T73 in the CRISPR LRRK2-KO clone, as expected (**Fig. 1A**). However, IF analysis of LRRK2 using c41-2 showed no difference in LRRK2 staining in PFA- or methanol-fixed LRRK2-WT and LRRK2-deficient MODE-K cells (**Suppl. Fig. 2A**). Among several tested LRRK2 antibodies, N241A/34 displayed the highest specificity for LRRK2 detection in MODE-K cells, particularly when combined with methanol fixation (**Suppl. Fig. 2B**). However, IF staining of mouse ileum with the N241/34 antibody was still unspecific, since it generated equally strong signals in LRRK2-KO tissues (**Suppl. Fig. 3**).

**Figure 1.**
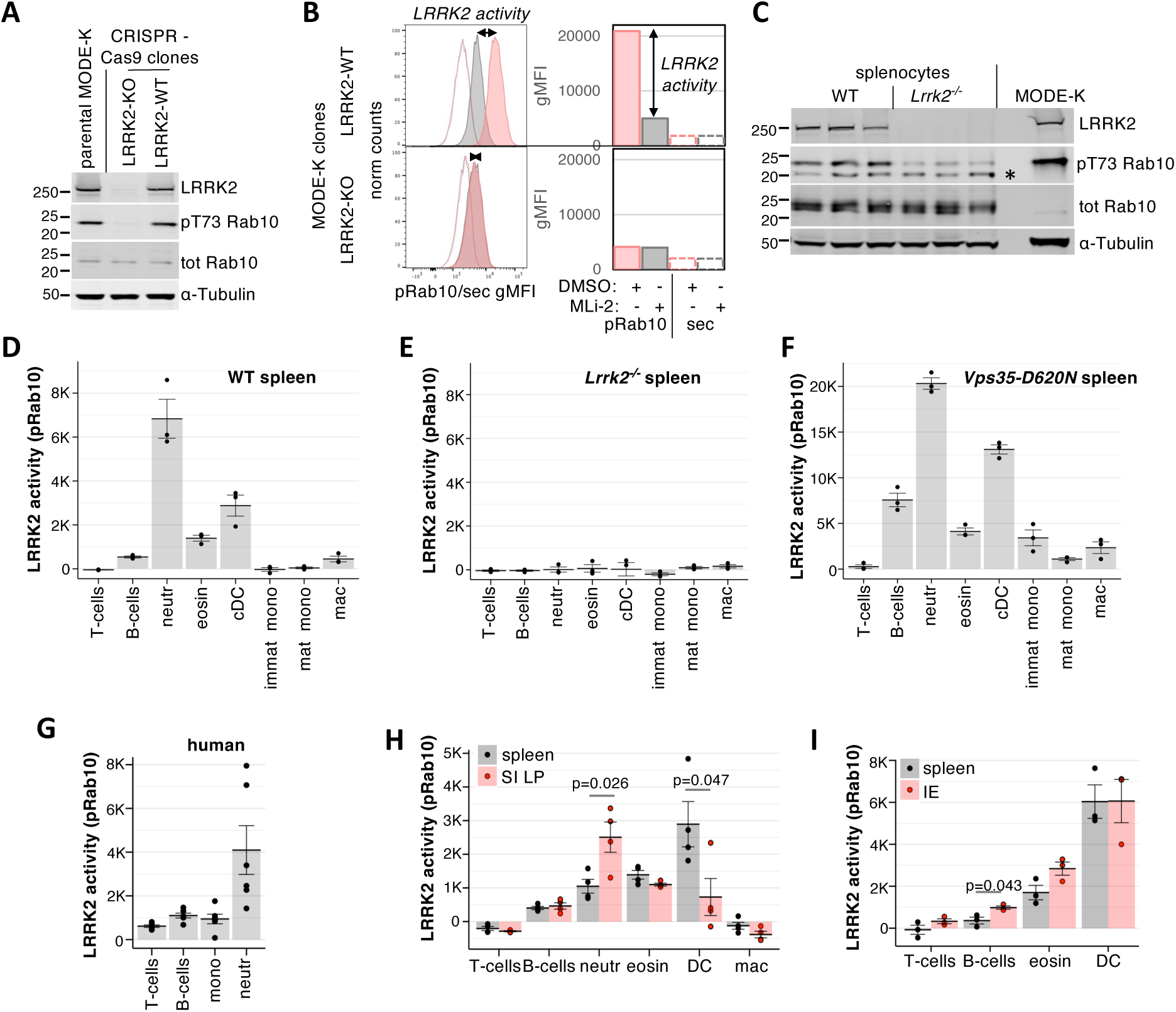
LRRK2 kinase activity is differentially regulated in splenic and intestinal immune cells. **A.** Loss of LRRK2 and phosphorylation of T73 on Rab10 in LRRK2-deficient (LRRK2-KO, second lane) CRISPR-Cas9-generated MODE-K clone. Parental MODE-K cells (left lane) and LRRK2-positive MODE-K clone (right lane) are shown as controls. Cell lysates were immunoblotted for total LRRK2 [N241A/34], pT73-Rab10 [MJF-R21] and total Rab10, with α-tubulin as a loading control. **B.** Flow cytometry of LRRK2-positive (top) or LRRK2-negative (bottom) MODE-K clones stained with anti-pT73 Rab10 ab [MJF-R21-22-5] followed by DyLight 649-labelled secondary antibody (solid lines), or with secondary antibody only (dashed lines), after 100 min incubation with 100 nM MLi-2 (grey) or vehicle (DMSO, pink). Mode-normalised histograms of fluorescence intensities are shown on the left, with geometric means of fluorescence intensities (gMFI) from the same data plotted on the right. The span of the arrows indicates LRRK2-dependent pRab10 signal, referred further as LRRK2 activity (pRab10). Note that this value is near-zero for LRRK2-deficient MODE-K clone. **C.** Isolated wild type (WT) and LRRK2-deficient (*Lrrk2^−/−^*) littermate splenocytes immunoblotted for total LRRK2 [N241A/34], pT73 Rab10 [MJF-R21], total Rab10 and α-tubulin as a loading control, to confirm the loss of LRRK2 in *Lrrk2^−/−^* cells. MODE-K cell lysate is included for comparison. A non-specific band in the pRab10 blot is indicated by a *. **D-E.** LRRK2 activity in WT (**D**) and *Lrrk2^−/−^* (**E**) splenocytes (n=3 mice each) shown in C. Splenocytes were treated with 100 nM of MLi-2 or 0.1% DMSO and stained with surface markers for immune cell types, including T-cells, B-cells, neutrophils (neutr), eosinophils (eosin), conventional dendritic cells (cDC), immature monocytes (immat mono), mature monocytes (mat mono) and macrophages (mac), before staining for intracellular pRab10. See Suppl. Fig. 4A for gating strategy. **F.** LRRK2 activity in splenocytes from mice (n=3) carrying VPS35-D620N mutation. **G.** Human peripheral blood mononuclear cells and neutrophils were isolated and stained for LRRK2-dependent pRab10. **H.** LRRK2 activity in indicated cell types isolated from spleen (grey) or small intestine lamina propria (SI LP, red) from (n=4) WT mice. See Suppl. Fig. 4B for gating. **I.** LRRK2 activity in indicated types of splenocytes (grey) or leukocytes of the small intestinal epithelium (IE) from (n=4) WT mice. D-I Dots depict LRRK2 activity (K = x1000) in individual mice or human donors, with means shown as bars. Error bars indicate SEM.

As an alternative approach, we explored flow cytometry-based detection. We tested several anti-LRRK2 antibodies and antibodies against phosphorylated LRRK2 substrates and selected an antibody against pThr73-Rab10, clone MJF-R21-22-5 (Lis *et al*, 2018), that showed the best specificity. Rab10 is the most highly phosphorylated Rab substrate of LRRK2 in most mouse tissues except the brain (Nirujogi *et al*, 2021). To ensure that the signal we observed was LRRK2-dependent, we included a control in which cells were treated with MLi-2, a specific LRRK2 kinase inhibitor (Fell *et al*, 2015). The pRab10 signal in both MLi-2 and vehicle treated cells was above signal generated by the secondary antibodies in the absence of primary antibodies (**Fig. 1B**). Importantly, in wildtype (WT) MODE-K cells, a sizeable increase in pRab10 fluorescence intensity was observed in vehicle-compared to MLi-2 treated cells. In contrast, in LRRK2-KO MODE-K cells, the pRab10 fluorescence intensity was low and unaffected by MLi-2 treatment (**Fig. 1B**). We defined specific LRRK2 activity in each cell as the difference between pRab10 geometric mean fluorescence intensity (gMFI) in the presence and absence of MLi-2:

LRRK2 activity = pRab10 gMFI_DMSO_ – pRab10 gMFI_MLi-2_

Thus, we established an assay that measures overall Rab10-directed LRRK2 activity in individual cells.

We next optimised this method to simultaneously measure LRRK2 activity in a complex mixture of cells isolated from tissues, using mouse spleen as a model organ. As a control, we used spleens obtained from LRRK2-KO mice (**Fig. 1C**). MLi-2/vehicle treatment and pRab10 staining was combined with staining for surface markers that identified a range of splenic cell types (**Suppl. Fig. 4A**). Our staining revealed a distinct pattern of LRRK2 activity, with highest level in neutrophils and conventional dendritic cells (cDC), followed by that in eosinophils, B-cells and macrophages, very low activity in monocytes and none in T-cells (**Fig. 1D**).

As expected, no LRRK2 activity was detected in splenocytes obtained from LRRK2-deficient mice (**Fig. 1E**), confirming the specificity of the assay. In contrast, splenocytes obtained from mice carrying the PD-associated VPS35 D620N mutation known to enhance LRRK2 activity (Mir *et al*, 2018) displayed strongly elevated LRRK2 activity in all cell types apart from T-cells (**Fig. 1F**). These studies validated the flow cytometric assay for assessing LRRK2 activity at a single cell level and defined the relative specific activity of LRRK2 in different immune mouse cells.

We also tested the assay in human peripheral blood mononuclear cells (PBMC) and isolated blood neutrophils, where we have previously shown high levels of LRRK2 activity (Fan *et al*, 2018). Despite the very low expression levels of LRRK2 found in human B-cells compared to monocytes (Fan *et al*., 2018), we saw comparable low levels of LRRK2-dependent pRab10 signal in B-cells and monocytes (**Fig. 1G**)(Fan *et al*., 2018), Interestingly, very low but measurable LRRK2 activity was also seen in PBMC T-cells. As expected, we saw high levels of LRRK2 activity in human neutrophils. Thus, in both human blood and murine spleens, neutrophils displayed the highest and T-cells the lowest level of LRRK2 signalling.

### 2. LRRK2 activity is regulated by tissue environment

We expanded our measurements to other mouse tissues where LRRK2 may play a role. Small intestinal lamina propria (SI LP) contains similar cell types as found in spleen. Interestingly, we found that LRRK2 activity was about 2.5-fold higher in SI LP neutrophils than in neutrophils obtained from spleen of the same mice, whereas dendritic cells from the SI LP possessed ∼4-fold lower LRRK2 activity (**Fig 1H**). In WT mice, LRRK2 activity was higher in several types of leukocytes that reside within small intestinal epithelium (IE) than in their counterparts obtained from spleen, with B-cells showing statistically significant increase in pRab10-directed LRRK2 activity (**Fig 1I**). Thus, LRRK2 activity in immune cells varies depending on the tissue context.

### 3. A novel EGFP-Lrrk2-KI reporter mouse reveals tissue- and cell-type specific expression of LRRK2 expression

Due to the challenges of assessing LRRK2 expression by IF discussed above, we developed an EGFP-Lrrk2 knock-in mouse line, in which the endogenous LRRK2 is N-terminally tagged with EGFP (**Fig. 2A**). Immunoblotting of LRRK2, GFP and pRab10 levels in WT, heterozygous (HET) and homozygous (HOM) mouse EGFP-Lrrk2-KI brain, spleen and large intestine lysates confirmed the presence of higher molecular weight LRRK2 protein that co-migrated with GFP signal in HET and HOM (**Suppl. Fig. 5A-C**). We noted that the addition of the of EGFP tag reduced total LRRK2 levels ∼50% in brain (**Suppl. Fig. 5A**), 33% in spleen (**Suppl. Fig. 5B**), without markedly affecting expression in intestine (**Suppl Fig 5C**). We also observed that pRab10 and pRab12 phosphorylation was reduced between ∼20% to ∼40% in HOM brain, spleen and intestine compared to WT (**Suppl. Fig. 5A-C**). As expected, an acute 2h administration of MLi-2 to mice markedly reduced pRab10 and pRab12 phosphorylation in all WT, HET and HOM tissues studied without impacting total levels of LRRK2 (**Suppl Fig 5A-C**). We also detected up to 50% reduction in LRRK2 activity, measured by flow cytometry as above, particularly in dendritic cells obtained from EGFP-Lrrk2-KI mice, compared to their WT littermates (**Suppl. Fig 5D**). These results indicate that the inclusion of a GFP tag at the N-terminus of LRRK2 moderately impacts LRRK2 expression and activity in most tissues and cells analysed. Note that no cleaved GFP was detected in GFP pulldowns (GFP-IP) from mouse lung tissue, indicating that the GFP signal corresponds to LRRK2 expression (**Suppl. Fig 5E**). Therefore, the EGFP fluorescence could serve as a readout to assess the relative levels of LRRK2 expression in different immune cells for the first time.

**Figure 2.**
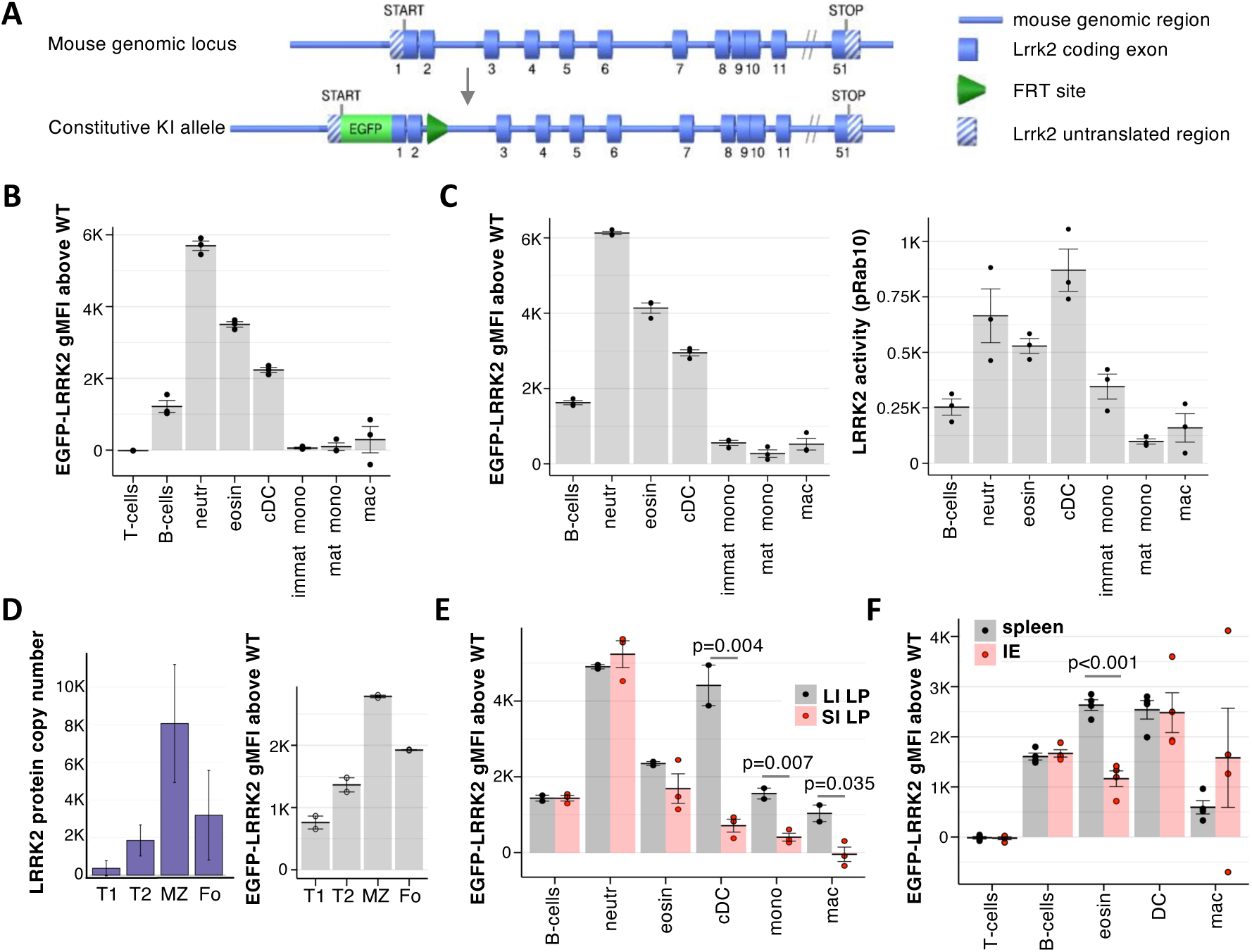
Measuring LRRK2 expression using EGFP-Lrrk2-KI reporter mouse model. **A.** Schematic of EGFP-Lrrk2-KI targeting. EGFP is inserted between N-terminal UTR and the start of the ORF of Lrrk2, in the context of endogenous Lrrk2 gene. **B.** EGFP fluorescence was measured by spectral flow cytometry in homozygous EGFP-Lrrk2-KI (n=3) splenocytes stained with cell markers and gated as in Suppl Fig 4, and EGFP-LRRK2 gMFI above background was calculated by subtracting the mean autofluorescence of the respective cell type measured in WT littermate mice (n=3). **C.** EGFP-LRRK2 expression (left) and LRRK2 activity (right) was measured in the same three EGFP-Lrrk2-KI mice as in B and Fig 1D. Three WT littermates were used to determine EGFP-LRRK2 fluorescence background. **D.** LRRK2 protein copy numbers (left) and EGFP-LRRK2 level (right) in splenic transitional 1 (T1), transitional 2 (T2), marginal zone (MZ) and follicular (Fo) B-cell subsets gated as in Suppl. Fig 6. Median LRRK2 copy numbers were quantified by label-free mass-spectrometry from WT splenocytes sorted into indicated subsets (Salerno et al, 2023). Data accessed from Immunological Proteome Resource, immpres.co.uk. **E.** EGFP-LRRK2 fluorescence in indicated lamina propria cell types isolated from large (LI LP, grey) and small (SI LP, red) intestine measured and displayed as in B. **F.** EGFP-LRRK2 fluorescence in splenocytes (spleen, grey) and cells obtained from small intestinal epithelium (IE, red) measured and displayed as in B. B-F Dots show values in individual mice, with means and SEM shown as bars and error bars. Statistically significant differences calculated by one-way ANOVA are indicated with p-values.

Among splenic immune cells, we found that the EGFP-LRRK2 expression was highest in neutrophils, followed by eosinophils, cDC and B-cells, with very low expression in macrophages and monocytes. EGFP-LRRK2 signal was undetectable in T-cells (**Fig. 2B**). The pattern of EGFP-LRRK2 levels was in some respects different from the pattern of LRRK2 activity measured in splenocytes of the same mice (**Fig. 2C**), indicating that LRRK2 activity is not determined solely by LRRK2 expression. To further validate the use of GFP flow cytometry in EGFP-Lrrk2-KI mouse-derived cells as a measure of LRRK2 expression, we analysed EGFP-LRRK2 expression in four major subpopulations of splenic B-cells, (transitional T1 and T2, follicular (Fo) and marginal zone (MZ), and compared it with publicly available mass-spectrometry data of LRRK2 expression in the same subsets of splenic B-cells (Immunological Proteome Resource, http://immpres.co.uk/, data from (Salerno *et al*, 2023). EGFP-LRRK2 expression (**Fig. 2D, right**) in all 4 subsets closely resembled the LRRK2 protein level measured by mass-spectrometry in the same B-cell subsets (**Fig. 2D, left**), orthogonally validating the EGFP flow cytometric signal as a readout for cellular LRRK2 expression.

We next asked how EGFP-LRRK2 expression in immune cells is affected by the intestinal environment. Intestinal eosinophils expressed less EGFP-LRRK2 than splenic eosinophils (**Fig. 2E**, compared with Fig. 2B). There was also a clear difference in EGFP-LRRK2 level between large and small intestinal lamina propria dendritic cells, monocytes and macrophages (**Fig. 2E**), supporting our earlier observation that tissue environment affects LRRK2. Furthermore, when several immune cell types isolated from intestinal epithelium (IE) were compared to their counterparts isolated from spleen, intestinal eosinophils showed significantly lower EGFP-LRRK2 expression, while EGFP-LRRK2 expression in T-cells, B-cells and dendritic cells were remarkably similar between these two tissues (**Fig. 2F**). These data support the conclusion that the EGFP-Lrrk2-KI mouse model serves as a reliable reporter of cellular LRRK2 protein expression and revealed tissue-specific and cell-type specific regulation of LRRK2 expression.

### 4. Inflammation alters LRRK2 activity and expression

To understand how inflammation affects LRRK2 in immune cells, we isolated splenocytes 24h after intraperitoneal injection of anti-CD3 antibodies that activates T-cells to release cytokines, driving transient inflammation (Swamy *et al*, 2015; Xu *et al*, 2021; Yaguchi *et al*, 2004; Zhou *et al*, 2004). This treatment induced inflammation apparent by significant change in cellular composition in the spleen (**Suppl. Fig. 7A**), as exemplified by a ∼35-fold increase in immature neutrophils (Deniset *et al*, 2017) (**Suppl. Fig. 7A, B**). Interestingly, anti-CD3-induced inflammation significantly increased LRRK2 activity in B-cells, neutrophils and monocytes (**Fig. 3A**). Measuring changes in EGFP-LRRK2 expression revealed a similar but not identical picture (**Fig. 3B**): the LRRK2 protein level was increased in B-cells, immature neutrophils and immature monocytes, while in contrast to its activity, LRRK2 protein was significantly reduced in mature neutrophils and cDC. The increase in EGFP-LRRK2 fluorescence in B-cells was not due to an increase in cell size associated with inflammation, since it was much larger than the anti-CD3 ab-induced increase in autofluorescence in WT samples (**Suppl. Fig. 7C**). Immunoblotting confirmed increases in both LRRK2 level and kinase activity in inflamed splenocytes, as determined by immunoblotting for total LRRK2 and Rab10 phosphorylation (**Fig. 3C**). These data emphasize that LRRK2 kinase activity and expression are decoupled in specific cell types during an inflammatory response.

**Figure 3.**
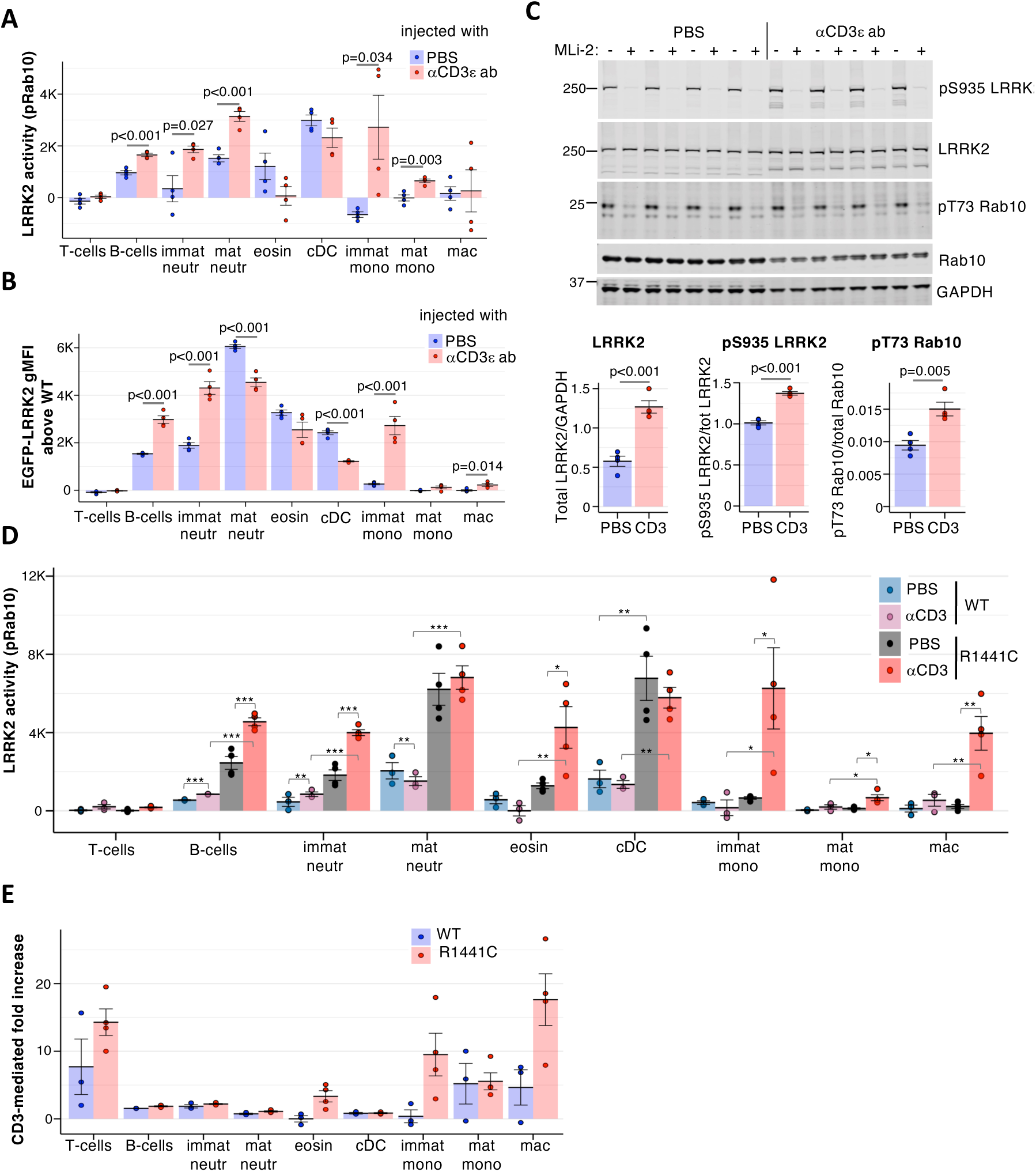
LRRK2 expression level and activity is modulated in a cell-type dependent manner by inflammation. **A.** LRRK2 activity was measured and displayed as in Fig 1C in splenocytes isolated from mice 24 h after i.p. injection of 100 μg of anti-CD3 antibody (red, n = 4) or the same volume (100 μl) of PBS (blue, n = 4). Significant differences between PBS- and anti-CD3-injected conditions are shown by p-values calculated by one-way ANOVA. **B.** EGFP-LRRK2 fluorescence in splenocytes from EGFP-Lrrk2-KI mice 24 h after i.p. injection of 100 μg of anti-CD3 (red, n= 4) or PBS (blue, n=4) co-stained with cell marker antibodies. The average autofluorescence from identically isolated, measured and analysed cells from PBS or anti-CD3 injected WT mice was subtracted as a background. Dots show data from individual mice, with means and SEM indicated by bars and error bars. Statistically significant differences are marked by p-values calculated by one-way ANOVA. **C.** Splenocytes from mice injected with either 100 μg anti-CD3 antibody or PBS shown in (A) were incubated for 30 min with media containing 100 nM MLi-2 (+) or equal volume of DMSO (−) and cell extracts immunoblotted for total LRRK2, pSer935 LRRK2, total Rab10 and pThr73 Rab10, with GAPDH as loading control. Quantifications for DMSO-treated samples are shown underneath, with p-values determined by 2-tailed t test. **D.** LRRK2 activity was measured as in (A) in splenocytes isolated from homozygous *Lrrk2-R1441C* mutant mice (R1441C) and their wild-type littermates (WT) 24 h after ip injection of 100 µg of anti-CD3ε antibody (red, n = 4 and 4) or the same volume (100 μl) of PBS (blue, n = 4 and 4). Statistically significant differences are indicated as *(0.01 < p < 0.05), **(0.001 < p < 0.01) and ***(p < 0.001) as measured by two-way ANOVA. **E.** Inflammation-induced change in LRRK2 activity in WT (blue) or *Lrrk2-R1441C* (red) splenic cells calculated as ratio between LRRK2 activities in anti-CD3-injected and the average pRab10 in PBS-injected mice of the same genotype from data shown in D. The bars show means of individual measurements shown as dots, with error bar indicating SEM. The contribution of genotype, inflammation and interaction between genotype and inflammation to LRRK2 activity determined by ANCOVA are shown in Table 2.

### 5. PD-associated mutation increases basal LRRK2 activity in mouse splenocytes and its activation by inflammation

We next investigated how LRRK2 activity in the context of anti-CD3-induced inflammation is affected by pathogenic LRRK2 mutations. For this, we exploited knock-in mice expressing the pathogenic LRRK2 R1441C mutation found in PD patients that has been previously shown to increase LRRK2 kinase activity in both murine cells and in humans (Alessi & Sammler, 2018). In both WT mice and HOM mice carrying LRRK2-R1441C mutation, anti-CD3 injection caused similar weight loss (**Suppl. Fig. 7D**) and increased mRNA levels of *Tnf, S100a8* and the IFN-responsive gene *Usp18* in the gut (**Suppl. Fig. 7E**). Moreover, cellular composition of the spleen did not show any significant changes between WT and mutant mice (**Suppl. Fig. 7F**), indicating that the overall level of inflammation was comparable in WT and *Lrrk2-R1441C* mice 24h post-injection with anti-CD3. Consistent with previous data (Iannotta *et al*, 2020), R1441C mutation markedly increased LRRK2 kinase activity in most splenic cell types in PBS-injected mice that served as a baseline control in this experiment (**Fig. 3D**, compare blue and grey bars). We found that anti-CD3-mediated inflammation further increased LRRK2-R1441C activity in most cell types (except T-cells, cDC and mature neutrophils), suggesting that the LRRK2-R1441C mutation and inflammation have different impacts on LRRK2 activity, either in parallel or in synergy (**Fig. 3D**, compare grey and red bars). Importantly, such response to inflammation was significantly exacerbated in B-cells, eosinophils and macrophages from *Lrrk2-R1441C* mice (**Fig. 3E**). Analysis of covariance (ANCOVA) confirmed a significant contribution of the interaction between the genotype and treatment in these cells (**Suppl. Table 1**). Thus, R1441C mutation not only enhances baseline LRRK2 activity, but also promotes its activation in the context of inflammation in some, but not all, cell types.

### 6. In vitro stimulation of splenocytes with anti-CD3 ab induces LRRK2 activity and EGFP-LRRK2 expression in B-cells

We next focused on inflammatory signal(s) that cause changes in LRRK2 expression and activity and asked whether anti-CD3-induced effects could be recapitulated *in vitro,* by culturing total splenocytes with anti-CD3/anti-CD28 antibodies to activate T-cells. Such stimulation resulted in marked enrichment of B-cells in the mixed splenocyte culture, and the loss of most other LRRK2-expressing cell types (**Suppl. Fig. 8A**); therefore, we focused on B-cells. We found that within mixed cultures, T-cell activation increased LRRK2 activity ∼2-fold in splenic B-cells (**Fig. 4A**) and EGFP-LRRK2 expression by ∼1.5-fold (**Fig. 4B**). B-cells isolated from such cultures (purity >95%, **Suppl. Fig. 8B**) displayed increased expression of MHCII and CD69 (**Fig. 4C**) compared to B-cells isolated from control cultures containing IL-7 only, suggesting that T-cell activation *in vitro* caused an activation of co-cultured B-cells. This activation was accompanied by a 3-fold increase in Lrrk2 mRNA level in B-cells from anti-CD3/CD28 stimulated cultures (**Fig. 4D**), indicating that LRRK2 is likely upregulated at the transcriptional level in these cells. Immunoblotting confirmed significant activation of LRRK2 phosphorylation of Rab10 in B-cells isolated from anti-CD3/CD28-stimulated cultures (**Fig. 4E**). Remarkably, EGFP-LRRK2 was also upregulated just by incubating isolated splenic B-cells with cell-free conditioned media collected from anti-CD3/CD28-activated splenocytes (**Fig. 4G**, see **Fig. 4F** for experimental layout). Incubation of B-cells isolated from *EGFP-Lrrk2*-KI mice with conditioned media collected from stimulated splenocytes led to activation of the B-cells (**Fig 4G**) and ∼2-fold increase in EGFP-LRRK2 expression (**Fig. 4H**), suggesting that LRRK2 is induced in B-cells by a soluble factor produced by activated T-cells. Thus, soluble factors produced by activated T cells can in turn activate B-cell expression and activity of LRRK2.

**Figure 4.**
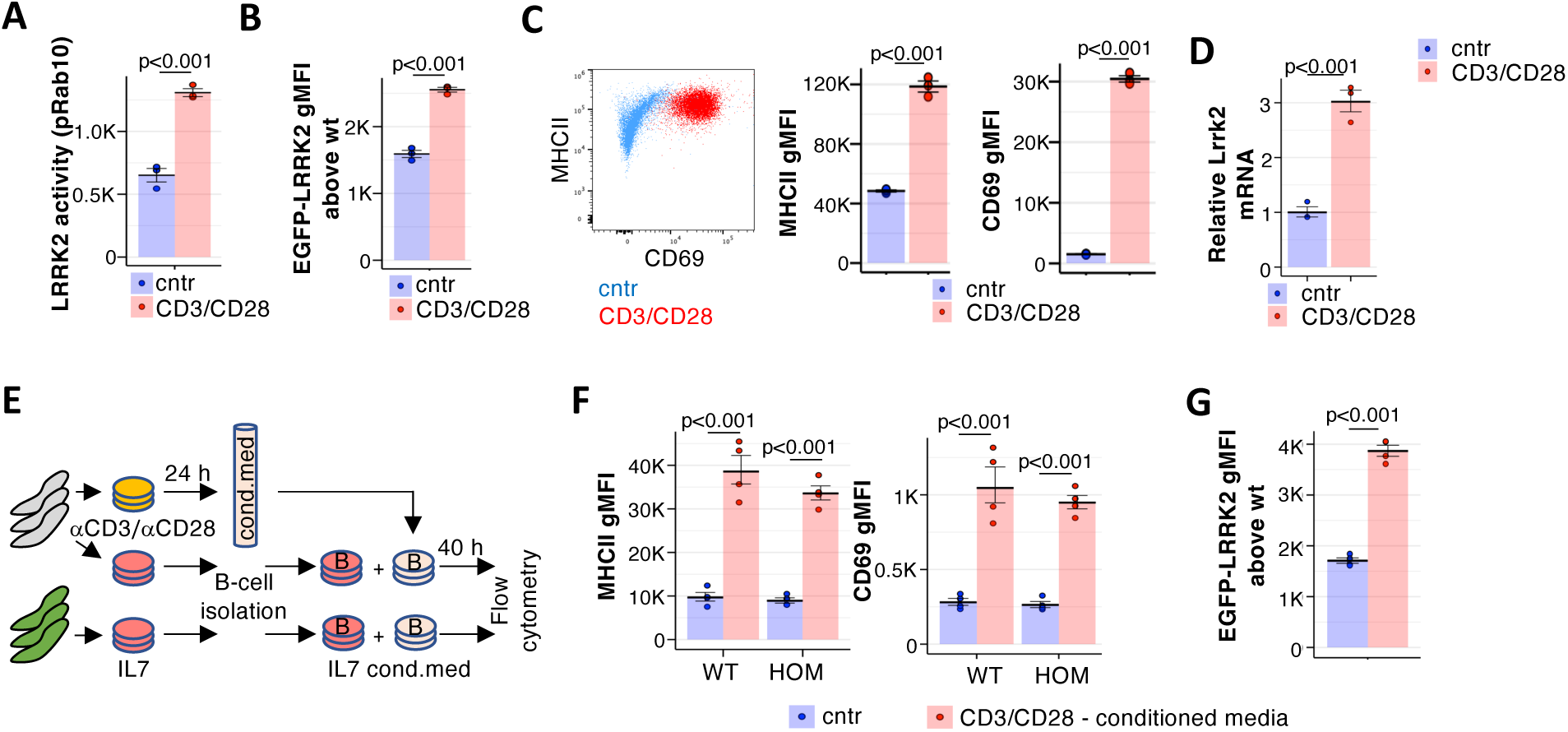
T-cell-dependent i*n vitro* stimulation enhances LRRK2 expression and activity in B-cells. **A.** Splenocytes from WT mice (n=3) were cultured for 24h with immobilised anti-CD3 and anti-CD28 antibodies (red), or with 1 ng/ml IL7 as a control (blue), and LRRK2 activity measured. **B.** Splenocytes from EGFP-Lrrk2-KI mice and their WT littermates (n = 3 each) were cultured as in A and stained with cell-type specific markers. EGFP-LRRK2 fluorescence was measured in B-cells from EGFP-Lrrk2-KI and WT mice, and autofluorescence of WT B cells was subtracted to determine EGFP-LRRK2 gMFI above WT background. **C**. B-cells were isolated from wild-type mouse splenocytes stimulated *in vitro* as in A, and their levels of MHCII and CD69 were measured. A representative scatterplot overlay of CD69 and MHCII in B-cells isolated from control (blue) or anti-CD3e/anti-CD28-induced (red) cultures is shown, along with the quantification of MHCII and CD69 in B-cells from control (blue) or induced (red) cultures. **D.** Relative mRNA levels of *Lrrk2* measured by qPCR in WT B-cells (n=3) from 24 h control (blue) or CD3/CD28-stimulated (red) splenocyte cultures. Bars are geometric means of fold change relative to a control group, with *Tbp* used as a reference gene. **E.** Experiment layout: splenocytes from EGFP-Lrrk2-KI or WT mice were cultured i*n vitro* in the presence of IL7 to support survival of resting cells or (for WT splenocytes) in the presence of immobilised anti-CD3e and anti-CD28 antibodies to stimulate T-cells. 24 h later, B-cells were isolated from resting WT and EGFP-Lrrk2-KI cultures and re-incubated for 40 h with IL7-supplemented media (IL7) or with conditioned media collected from stimulated (CD3/CD28) WT cells before cells were harvested and measured. **F**. Activation of B-cells at the end of the 40 h incubation with CD3/CD28-conditioned media (red), compared with IL7-only treated cells (cntr) as outlined is apparent from significant increase in MHCII (left panel) and CD69 (middle panel) for both genotypes (either wild type [WT] or EGFP-Lrrk2-KI homozygous [HOM]).**G.** EGFP-LRRK2 expression in isolated B-cells incubated with either IL7 control (blue) or with CD3-CD28-conditioned media (red) was measured and displayed as in B. A-G Measurements from individual mice are shown as dots, with mean values and SEM depicted as bars and error bars. Statistical significance was calculated using ANOVA, and p-values indicated above each plot.

### 7. IL-4 induces LRRK2 activation in B-cells

To identify the soluble factor(s) enhancing LRRK2 expression, we tested the ability of neutralizing antibodies against several cytokines likely to be produced by activated T-cells and present in anti-CD3/CD28 stimulated conditioned media to block LRRK2 induction. We found that antibodies against IL-4, but not IFNγ, TNF or IL-17A, significantly inhibited EGFP-LRRK2 induction by conditioned media (**Fig. 5A**). Furthermore, recombinant mouse IL-4 was able to upregulate EGFP-LRRK2 expression and increase LRRK2-dependent Rab10 phosphorylation in B-cells (**Fig. 5B-C**). The effect of IL-4 was much more pronounced than the effect of IFNγ known to upregulate LRRK2 in human monocytes, human neurons and murine macrophages (Cook *et al*., 2017; Gardet *et al*., 2010; Panagiotakopoulou *et al*, 2020). In our system, IFNγ induced only a small increase in the level and activity of LRRK2 of B-cells and was not responsible for LRRK2 upregulation by anti-CD3/CD28 conditioned media. IL-4 also induced LRRK2 at the transcriptional level (**Fig. 5D**). Furthermore, quantitative mass spectrometric analysis of LRRK2 expression in IL-4 treated B-cells showed a ∼2.5-fold increase in LRRK2 copy numbers ((**Fig. 5E**). To determine whether the IL-4-dependent induction of LRRK2 expression was a result of population shift towards a subset with a higher expression of LRRK2 (such as Fo or MZ, see Fig 2D), we combined EGFP-LRRK2 measurements with B-cell subset analysis. Culturing purified B-cells in homeostatic levels of IL-7 (control) for 24h resulted in loss of MZ, and addition of IL-4 further shifted the relative abundance of B-cells towards transitional CD93-high T2 state, with a concomitant reduction in T1 and Fo B-cells (**Fig. 5F, left**). Regardless, we observed that IL-4 significantly upregulated EGFP-LRRK2 in all detectable B-cell subsets (**Fig. 5F, right**), indicating that the IL-4-dependent induction of LRRK2 is unrelated to B-cell maturation state. Finally, immunoblotting of LRRK2, pRab10 and total Rab10, in purified mature B cells cultured with IL-4 also showed that total LRRK2 levels and LRRK2 activity were significantly increased by IL-4 (**Fig. 5G**). Thus, we identify IL-4 as a novel inducer of LRRK2 expression and kinase activity in B-cells.

**Figure 5.**
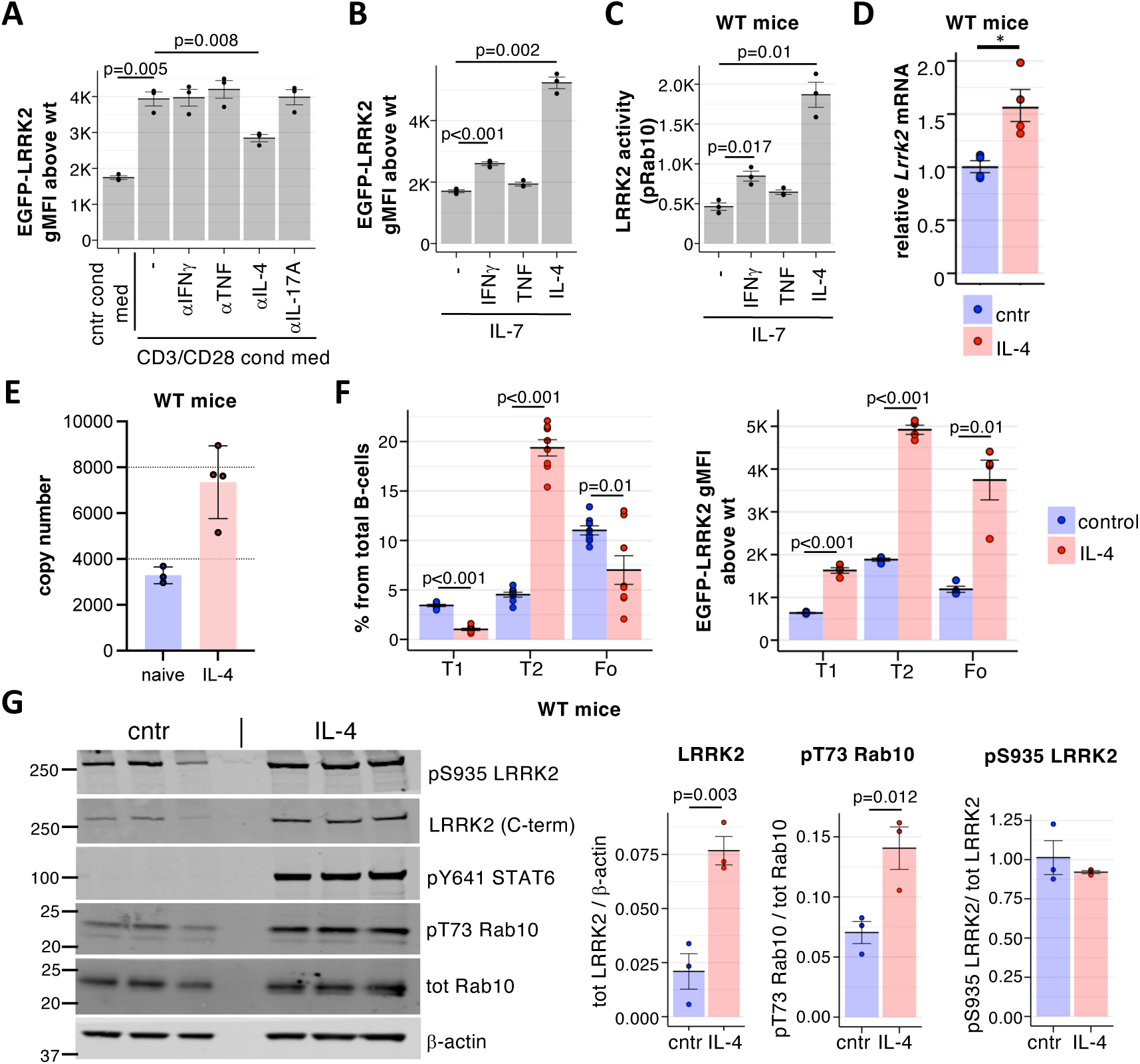
IL-4 induces LRRK2 expression and activity in splenic B-cells. **A.** EGFP-LRRK2 fluorescence was measured in B-cells isolated from EGFP-Lrrk2-KI spleens (n = 3) and incubated for 40h with conditioned media from splenocytes as described in Fig. 4E, supplemented where indicated with 10 µg/ml of neutralising antibodies, and the autofluorescence of identically treated and measured B-cells from wild-type mice (n = 2) was subtracted as a background. p-values show statistically significant changes in EGFP-LRRK2 expression calculated by paired t-test. **B**. B-cells isolated from EGFP-Lrrk2-KI spleens were cultured for 40 h in media containing 5 ng/ml IL-7 with or without 10 ng/ml IFNγ, 100 ng/ml TNF or 10 ng/ml IL-4 as indicated, and EGFP fluorescence above background measured as in A. **C.** LRRK2 activity and activation markers in B-cells isolated from three WT mice treated as in B was measured. **D.** Relative mRNA levels of *Lrrk2* measured by qPCR in WT B-cells (n=3) from 24 h control (blue) or IL-4 stimulated (red) cultures. Bars are geometric means of fold change relative to a control group, with *Tbp* used as a reference gene. **E.** Graph shows the estimated copy number of LRRK2 per cell (estimated using the histone proteome ruler method) in lymph node B cells stimulated for 40 hours with 10ng/ml IL4. **F.** B-cells isolated from EGFP-Lrrk2-KI and WT mice and cultured for 40 h with (red) or without (blue) 10 ng/ml IL-4 as in B were co-stained with markers for splenic B-cell subsets, and subset frequencies (left) and EGFP-LRRK2 expression above background (right) were measured in T1, T2, and Fo B-cell subsets gated as in Suppl. Fig 6. Significance calculated by paired t-test. Dots show values in individual mice, with means and SEM shown as bars and error bars. **G.** Immunoblot of WT splenic B-cells isolated using positive selection and cultured for 24 h with (IL-4) or without (cntr) 10 ng/ml IL-4. Lysates from equivalent cell numbers were loaded and blotted for total LRRK2 [N241/34], pS935-LRRK2, pY641 STAT6, total Rab10 [605B11] and pT73-Rab10. β-actin was used as a loading control. Quantifications of the total LRRK2 protein levels, LRRK2 activity towards Rab10, pS935-LRRK2 and pSTAT6 are shown as bars (means ± SEM), with statistical significance calculated by paired t-test indicated by p-values.

## Discussion

The relatively low and highly variable prevalence of pathogenic LRRK2 mutations among PD patients (Simpson *et al*, 2022) prompted development of many assays to assess LRRK2 status for patient stratification (Rideout *et al*, 2020). Most such assays utilise immunoblots, ELISA, or proteomics to examine the total level of LRRK2, its phosphorylation, or the phosphorylation of its substrates. In this study, we have independently developed a flow cytometry-based method to quantify LRRK2 activity with single cell resolution, that builds upon previously published flow cytometry assays to assess LRRK2 kinase activity (Dhekne *et al*, 2023; Hakimi *et al*., 2011; Wallings *et al*, 2022). We introduced a crucial control, namely treating cells ± MLi-2 prior to flow cytometry. Since the background signal of pRab10 varies between different cell types, this control is essential for the specific evaluation of LRRK2-dependent phosphorylation of Rab10, allowing for a more accurate estimate of LRRK2 kinase activity across different cell types. Our approach allowed us to confidently measure LRRK2 activity in different types of immune cells, including T-cells, neutrophils, monocytes, macrophages that have been implicated in infection control and in Crohn’s disease. Importantly, we show for the first time that eosinophils also express active LRRK2 at levels comparable to B-cells and DCs. Since eosinophils have recently been implicated as key player in intestinal defence and colitis (Gurtner *et al*, 2023), it will be interesting to evaluate LRRK2 functions in these cells.

We complemented the pRab10 assay with the development of the EGFP-Lrrk2-KI reporter mouse. Although the reporter was initially designed as a fluorescent tracker for imaging LRRK2 localisation in cells and tissues, the low expression of LRRK2, combined with high and variable autofluorescence in intestinal tissue in particular precluded its use for microscopy. Even in neutrophils, which express highest level of LRRK2 among immune cells, there are less than 10,000 copies of LRRK2 per cell (Sollberger *et al*, 2024). However, the EGFP signal was sufficient for flow cytometry-based measurements, where background autofluorescence of each cell type was taken into account and subtracted.

We found that inflammation strongly enhances LRRK2 expression and activity in B-cells. While the role of B-cells in Crohn’s disease is not yet fully established (Zogorean & Wirtz, 2023), these cells are important players in several inflammatory autoimmune diseases including Systemic Lupus Erythematosis (SLE) and Multiple Sclerosis (MS). LRRK2 is upregulated in B-cells of patients with SLE (Zhang *et al*, 2019). Interestingly, the authors also found increased LRRK2 expression in published transcriptomic data (Liu *et al*, 2017b) from human CD19^hi^/FSC^hi^ B-cells produced *in vitro* by co-culturing B-cells with anti-CD3-CD28 stimulated CD4+ T-cells. These cells show activation and memory markers, have a higher propensity to produce IgG and IgM and are thought to represent the activated pathological CD19^hi^ B-cells subset found in the periphery of SLE patients (Liu *et al*., 2017b). Similar to these cells, the LRRK2-upregulating activation of B-cells that we observed was accompanied by elevated CD19, IgM, cell size and granularity and required activated T-cells for their development, so it is possible that we are dealing with the same type of cells. However, human CD19^hi^ B-cells relied on the direct interaction with stimulated T-cells for their development, since their numbers and ability to produce IgM and IgG was dramatically reduced by blocking ICAM-1 that mediates such interaction, CD40L or OX40 that are normally expressed on the surface of activated T-cells. In contrast, we found that supernatant from anti-CD3-CD28 stimulated splenocytes, or Th2-type cytokine IL-4 was sufficient to upregulate LRRK2 and induce similar activation phenotype in mouse splenic B-cells.

IL-4 is a pleiotropic cytokine that promotes B-cell maturation, IgG class switching and enhances production of IgG and IgA. Although it was initially described as a B-cell growth factor, on its own IL-4 is not able to promote proliferation of resting B-cells (Howard *et al*, 1982), but it is co-stimulatory together with LPS, CD40L and antigenic stimulus. IL-4 has been reported to induce autophagy in B-cells that in turn promotes B-cell survival and antigen presentation (Xia *et al*, 2018), and recent studies show LRRK2 is activated downstream of non-canonical autophagy / Conjugation of ATG8 to Single Membranes (CASM) pathway (Bentley-DeSousa & Ferguson, 2023; Eguchi *et al*, 2024). In addition, IL-4 is absolutely required for antigen-specific IgA production in the intestine after oral immunisation, and for germinal centre formation in Peyer’s patches in the murine small intestine (Vajdy *et al*, 1995). It is interesting to note that Lrrk2-KO mice have increased levels of serum IgA (Kubo *et al*, 2016). Whether IL-4 induced LRRK2 in B-cells is important for IgA switching in B-cells remains to be determined, but the link with autophagy suggest a potential pathway through which LRRK2 might function. Further, IL-4 can have both pro- and anti-inflammatory roles in disease, and in one mouse model of Crohn’s disease-like ileitis, a pathogenic role has been assigned to IL-4 (Bamias *et al*, 2005). Thus, this newly discovered IL-4-LRRK2 pathway in B-cells may also contribute to autoimmune diseases.

In conclusion, we describe the utility of a LRRK2 reporter mouse model and a flow cytometric based assay to robustly quantify LRRK2 kinase activity on a per cell basis. Using these assays we demonstrate inflammation-driven activation of LRRK2 in vivo that is highly regulated by both cell type and tissue context. These tools enable dissection of spatial and temporal activation of LRRK2 in disease context, advancing our understanding of the critical roles different cells play in the pathological functions of LRRK2 across diseases such as SLE, PD, Crohn’s, and others.

## Methods

### Cell lines

Murine small intestinal epithelial cell line MODE-K (Vidal *et al*, 1993) (kind gift from D. Kaiserlian, Lyon) was cultured under DMEM containing 10% Foetal Bovine Serum (FBS) and 100 U/ml penicillin-streptomycin. To generate LRRK2-KO MODE-K clones, parental MODE-K cells were co-transfected with the pX335 plasmid encoding spCas9n D10A nickase and an antisense guide GCAGCCCTGACAGGCGCCAC(TGG), and the pKN7 plasmid with puromycin resistance gene and a sense guide gCTCTGAAGAAGTTGATAGTC(AGG), both targeting exon 1 of mouse LRRK2, using Lipofectamine LTX with PLUS Reagent (Invitrogen), and selected for 2 days with 5 µg/ml puromycin. Single cells from the resulting culture were sorted into 96 well plates using MA900 Cell Sorter (Sony Biotechnology) and the colonies were expanded, collected and screened for the loss of LRRK2 and Rab10 phosphorylation using immunoblotting and pRab10 flow cytometry (see below).

### Mice

Mice were bred and maintained with approval by the University of Dundee ethical review committee under a UK Home Office project license (PP2719506) in compliance with UK Home Office Animals (Scientific Procedures) Act 1986 guidelines. Lrrk2-KO (*Lrrk2^−/−^*) mice (Parisiadou *et al*, 2009) were from Prof. Huaibin Cai (NIH, Bethesda) and maintained on C57BL/6J background. *Lrrk2-R1441C* knock-in mice (Tong et al, 2009) were obtained from the Jackson Laboratory (JAX strain #009347) and maintained on C57BL/6J background. *Vps35-D620N* knock-in mice (Mir et al., 2018) were from the Jackson Laboratory (strain #023409). All mice were maintained in a pathogen-free conditions in a standard barrier facility on a 12 h light/dark cycle at 21°C in individually ventilated cages with sizzler-nest material and fed an R&M3 diet (Special Diet Services, UK) and filtered water ad libitum. Cages were changed at least every 2 weeks.

### Generation of EGFP-Lrrk2 knock-in mice

EGFP-LRRK2 knock-in mice were custom-made by Taconic Biosciences. The targeting vector was constructed by inserting EGFP sequence between translation initiation codon and the second codon in exon 1 of LRRK2 (NCBI transcript NM_025730.3), and inserting a positive selection marker, PuroR, flanked by FRT sites into intron 2. The targeting vector was generated using BAC clones and transfected into C57BL/6N Tac ES cell line. After homologous recombinant clones were selected, the selection marker was removed by Flp-mediated excision, producing the constitutive KI allele encoding EGFP-LRRK2 fusion protein expressed from endogenous *Lrrk2* promoter. Mice were backcrossed at elast 10 generations to C57BL/6J.

### Anti-CD3-induced inflammation in vivo

Mice were intraperitoneally (ip) injected with 100 µg of Ultra-LEAF (Low Endotoxin, Azide-free) purified anti-mouse monoclonal anti-CD3ε antibody, clone 145-2C11 (Biolegend, #100340) in Dulbecco’s Phosphate Buffered Saline (PBS, Gibco) in 100 µl volume, or 100 µl of PBS and maintained for 24 h under observation before tissue harvesting.

### MLi-2 treatment of EGFP-Lrrk2 mice

For immunoblotting analysis of EGFP-Lrrk2 mice, tissues were collected from mice injected subcutaneously with either the LRRK2 specific inhibitor MLi-2 or vehicle, 40% (w/v) (2-Hydroxypropyl)-β-cyclodextrin. Mice were injected with MLi-2 at 30 mg/kg final dose or vehicle and euthanized by cervical dislocation 2 hours later. This dose and duration of treatment have been shown to effectively inhibit LRRK2 in all tissues examined including brain (Nirujogi *et al*., 2021).

### Tissue H&E and immunofluorescence

Isolated mouse small intestine was flushed with HBSS and 1.5 cm fragments of ileum were fixed in 4% Paraformaldehyde (PFA) in PBS (pH = 7.4) for 24 h at +4^0^C with gentle rocking, and washed three times in PBS prior paraffin perfusion and embedding into paraffin blocks. 5 µm sections from paraffin-embedded tissues were deparaffinised using Histo-Clear xylene substitute, rehydrated, and either stained in Mayer’s haemotoxylin & Eosin, or processed for immunofluorescence (IF). For IF, the antigen retrieval was performed in 10 mM Trisodium citrate containing 0.05% Tween 20, pH = 6.0, in a pressure cooker. The tissues were permeabilised for 20 min in 1% NP40 in PBS, blocked for 1 hour in PBS containing 2% Bovine Serum Albumin (BSA), 5% normal goat serum and 0.1% Triton X-100 with one drop of Mouse on Mouse Blocking Reagent (VectorLabs, #MKB-2213) per 600 µl, and stained with the primary antibodies diluted in PBS containing 2% BSA and 0.1% TritonX100 (anti-LRRK2, clone MJFF2 (c41-2), rabbit monoclonal antibody at 1:100 dilution and anti-E-cadherin (BD Bioscences, clone 36/E-cagherin (RUO), #610182) antibody at 1:150 dilution, or anti-LRRK2, clone N241A/34, mouse monoclonal antibody at 1:200 dilution) at 4^0^C overnight, followed by staining with appropriate secondary antibodies (goat anti-rabbit IgG Alexa Fluor 568, Invitrogen #A-11036, and goat anti-mouse IgG Alexa Fluor Plus 488, Invitrogen #A32723), both highly cross-adsorbed to avoid any cross-species detection) diluted to 1:500 in PBS containing 2% BSA and 0.1% TritonX100 for 1 hour at room temperature. Washing was done with PBS throughout the procedure. The tissues were counterstained with 1 µg/ml DAPI and slides were mounted in Vectashield Vibrance antifade mounting media (Vector Laboratories, H-1700). Fragments of the lung were fixed at room temperature in 2% PFA in PBS (pH = 7.4) for 2-3 h, washed three times in 50 mM ammonium chloride in PBS, rinsed and incubated in 30% sucrose in PBS for 24 h at +4^0^C, before embedding into OCT compound (Agar Scientific, AGR1180) on dry ice. 15 µm cryosections were permeabilised for 15 min in 0.2% saponin in PBS, blocked in PBS containing 2% BSA and 0.025% saponin and incubated with anti-LRRK2 [MJFF2 (c41-2)] antibody diluted 1:100 in PBS/2% BSA/0.025% saponin overnight at +4^0^C, followed by 1 h incubation with goat anti-rabbit AlexaFluor-568 secondary ab (Invitrogen, #A11036) at 1:500 dilution with Acti-stain^TM^ 488 phalloidin (Universal Biologicals, #PHDG1-A) at 1:75 dilution in PBS/2% BSA/0.025% saponin. Slides were counterstained with 1 µg/ml DAPI and mounted in Vectashield Vibrance antifade mounting media.

Stained tissues were imaged on confocal Zeiss 710 microscope operated by ZEN software (Zeiss) using a 20x Plan Apochromat 0.8 NA dry objective with zoom 0.6-1. All images, including controls in which primary antibodies were omitted, were acquired using identical microscopy settings. Resulting two- or three-channel images were further processed and assembled in OMERO.figure (https://www.openmicroscopy.org/omero/figure/), identically within each experiment.

### Cell immunofluorescence

Parental MODE-K cells and LRRK2-KO MODE-K clone were seeded onto autoclaved glass coverslips placed in the wells of 6-well plates, 1.6 × 10^5^ cells per well 24 h before fixing/permeabilising them with either 4% PFA in PBS supplemented with 1% Triton X-100 for 10 min at +37^0^C or with ice-cold 100 % methanol for 5 min at −20^0^C. Coverslips with cells were washed with PBS, blocked in PBS containing 2% BSA, 5% Normal Goat Serum and 0.1% Triton X-100 and stained overnight at +4^0^C in either N241A/34 or c41-2 anti-LRRK2 antibody, diluted 1:100 or 1:200, respectively, in above blocking buffer, followed by several PBS washes and 1 h incubation with secondary antibodies (either goat anti-rabbit IgG Alexa Fluor 568 or goat anti-mouse IgG Alexa Fluor Plus 488) diluted 1:500 in blocking buffer, before counterstaining with 1 µg/ml DAPI and mounting in ProLong Gold antifade reagent, Thermo Fisher Scientific, #P36930. Staining in which primary antibody were omitted were used as controls. For each antibody and appropriate secondary-only controls, cells were imaged on confocal Zeiss 710 microscope operated by ZEN software (Zeiss) using a 63x Plan Apochromat 1.4 NA oil objective using identical setting, and images were further processed and assembled in OMERO as above.

### Isolation of primary cells from mouse tissues

Splenocytes were obtained by mashing freshly isolated spleen through Falcon^TM^ 70 µm nylon Cell Strainer in PBS containing 2% Foetal Bovine Serum (FBS), followed by erythrocyte lysis using ACK Lysing Buffer (Gibco), resuspended in RPMI 1640/10%FBS and filtered through above 70 µm Cell Strainer. For B-cell isolation, splenocytes were washed in PBS and resuspended in PBS containing 2%FBS and 2 mM EDTA before negative selection using EasySep Mouse Pan-B Cell Isolation Kit (StemCell Technologies, #19844A) following manufacturer instructions. Alternatively, where indicated, B-cells were isolated using positive selection on Mojosort Streptavidin Nanobeads (Biolegend, #480016) coupled with Biotinylated anti-CD23 antibodies (BD Pharmingen, BD biosciences, #553137), as follows: splenocytes were resuspended in FACS buffer (PBS supplemented with 2%FBS and 2 mM EDTA) up to 1×10^8^ cells/ml and incubated with anti-CD23 antibody at 1:200 dilution for 10 min at +4^0^C, spun down and resuspended in the fresh FACS buffer and the Mojosort Streptavidin Nanobeads, 10 beads µl per 100 µl of cell suspension containing 10^7^ cells. After 15 min incubation on ice, cells with beads were washed once in FACS buffer, resuspended in 2.5 ml of FACS buffer and isolated on an EasySep magnet (Stemcell Technologies, #18000). The magnet-bound fraction was collected, and isolation was repeated twice more to increase purity. To obtain small intestine or large intestine lamina propria (SI LP or LI LP) cells, small intestine or large intestine was excised, flushed with Hanks’ Balanced Salt Solution (HBSS, Gibco) and cut longitudinally and across into ∼1 cm pieces that were washed by vigorous shaking in PBS three times, collecting tissues in the 70 µm Cell Strainers between the washes, before incubating for 30 min at 37^0^C in a bacterial shaker at 225 rpm in a strip buffer (PBS containing 5% FBS, 1 mM Dithiothreitol (DTT) and 1 mM EDTA), replacing the strip buffer with the fresh one 10 min after the start. Tissues were then washed three times by vigorous shaking in PBS and incubated for 45 min at 37^0^C in a bacterial shaker in 10 ml of Digest buffer (RPMI 1640 containing 10% FBS, 100 U/ml penicillin-streptomycin, 2mM L-glutamine and 1 mg/ml collagenase/dispase (Roche, #11097113001) and 20 µg/ml Recombinant RNase-free DNase I (Roche, #04716728001)). Digested cell suspension was filtered through 70 µm Cell Strainer and washed twice in RPMI 1640 containing 10%FBS. Intraepithelial lymphocytes were obtained as described (James *et al*, 2020). Briefly, isolated small intestine was flushed with HBSS, cut as above into warm RPMI 1640 containing 10% FBS, 2mM L-glutamine, 100 U/ml penicillin-streptomycin and 1 mM DTT and incubated for 40 min at room temperature with continuous shaking. Tissue was spun down, resuspended in the same media without DTT and vortexed for 3 min before collecting the dissociated cells through 100 µm Cell Strainer, and the resuspension/vortexing/cell collection procedure was repeated two more times with remaining tissue, combining the IEL-containing flow through from all rounds. The cells in the flow through were then pelleted, resuspended in 40% Percoll, overlayed over 75% Percoll and spun for 30 min at 700g, with gentle acceleration and deceleration. Cells at the interface between 40% and 75% Percoll gradient were collected and washed once with RPMI 1640 containing 10% FBS, 2mM L-glutamine and penicillin-streptomycin.

### Cell treatment

When indicated, splenocytes were seeded in 6-well plates and incubated for 30 min with RPMI 1640 with 10% FBS containing either 100 nM MLi-2 or 0.1% DMSO. For *in vitro* anti-CD3/CD28 stimulation, isolated splenocytes were seeded in RPMI 1640 media containing 10% FBS, 100 U/ml penicillin-streptomycin, 2 mM L-glutamine, 1 mM sodium pyruvate, 1% non-essential amino acids, 50 µM β-mercaptoethanol, and 2.5 mM HEPES into 6-well plates pre-coated for more than 1 h with 5.4 µg of Ultra-LEAF^TM^ purified anti-mouse CD3e antibody (clone 145-2C11, Biolegend, # 100340) and 3 µg of Ultra-LEAF^TM^ purified anti-mouse CD28 antibody (clone 37.51, Biolegend, # 102116) per well, and incubated in standard cell culture conditions for 24 h before harvesting. As a control, cells were incubated in the above media supplemented with 1 ng/ml murine IL-7 (PeproTech, #217-17). Isolated B-cells were incubated for 40 h in the above media that was diluted 1:1 with filtered control or anti-CD3/CD28-stimulated conditioned media collected from splenocytes after 24 h incubation described above or supplemented with 5 ng/ml murine IL-7. Throughout the 40 h incubation, cells were treated with 10 µg/ml of Ultra-LEAF^TM^ purified anti-mouse IFNγ antibody (clone R4-6A2, Biolegend, #505709), 10 µg/ml of Ultra-LEAF^TM^ purified anti-mouse TNF antibody (clone MP6XT22, Biolegend, #100340), 10 µg/ml of Ultra-LEAF^TM^ purified anti-mouse IL-17A antibody (clone QA20A33, Biolegend, # 946504), 10 µg/ml of Ultra-LEAF^TM^ purified anti-mouse IL-4 antibody (clone 11B11, Biolegend, #504122), 10 ng/ml animal-free recombinant murine IFNγ (PeproTech, #AF-315-05), 100 ng/ml of recombinant murine TNF (PeproTech, # 315-01A) or 10 ng/ml of recombinant murine IL4 (PeproTech, #214-14) or left untreated as indicated.

### Human peripheral blood neutrophil and PBMC isolation

The research was approved by the East of Scotland Research Ethics Service (REC Reference 19/ES/0031) and informed consent was obtained from all participants. Neutrophils and peripheral blood mononuclear cells (PBMCs) were isolated from fresh human peripheral blood from healthy donors as described previously (Fan *et al*., 2018). Briefly, neutrophils were isolated from human whole blood based on immuno-magnetic negative isolation technique using the EasySep Direct Human Neutrophil Isolation Kit (STEMCELL Technologies, Cat# 19666) described in detail in (Fan *et al*, 2020). PBMCs were isolated from whole blood using density centrifugation with Ficoll-Paque PREMIUM density gradient (GE Healthcare, Cat# 17-5442-02) and SepMate™ tubes (Stemcell, Cat# 85450) as described in detail in (Sarhan *et al*, 2021). After isolation, neutrophils and PBMCs from each donor were resuspended in RPMI and treated with 200nM MLi-2 or DMSO (0.1% v/v). The cells were then subjected to the same flow cytometric workflow for pRab10 as described below.

### Flow cytometry

For EGFP-LRRK2 measurements and cell profiling, isolated cells were stained with viability dye (DAPI, LIVE/DEAD^TM^ Fixable Near-IR (Thermo Fisher Scientific, #L10119) or Ghost Dye Violet 450 (Cytek, Cat nr. 13-0863-T100)) followed by staining with the antibody mixture (see table below for details) for identifying splenocytes and lamina propria cells (against CD45, MHCII [I-A/I-E, CD11c, CD11b, Ly6G, B220, TCRb, SiglecF, CD64, Ly6C and F4/80); intraepithelial immune cells (against CD45, MHCII [I-A/I-E], CD11c, NK1.1, SiglecF, CD64, CD3 and CD19), B-cell subsets (against CD19, CD93, IgM, CD23, CD21/CD35) or for assessing B-cells purity and activation (CD19, B220, MHCII and CD69) and measured on 3-laser Cytek® Aurora spectral cytometer system operated by SpectroFlo® software (Cytek).

For measuring LRRK2 activity, cells were stained with LIVE/DEAD^TM^ Fixable Near-IR dye or Ghost Dye Violet 450 followed by 30 min incubation with either 200 nM MLi-2 (synthesised by Natalia Shapiro, University of Dundee) or equal amount of vehicle (0.2% DMSO) in PBS containing 2% FBS, 1 mM EDTA and FC block (Purified anti-mouse CD16/32, clone 93, BioLegend, Cat Nr101302) at 1:200 dilution at room temperature, before adding equal volume of desired antibody mixture (as for EGFP-LRRK2 measurements, except for intraepithelial immune cells it was against EpCAM, CD45, MHCII, CD11c, CD3 and CD19) and incubating for another 30 min. Cells were washed, fixed for 20 min with 2% PFA in PBS and washed/permeabilised in 1x Permeabilisation buffer (Invitrogen by Thermo Fisher Scientific, #00-8333-56), followed by overnight staining with anti-pT73 Rab10 antibody (clone MJF-R21-22-5, AbCam ab241060) diluted 1:500 in 1x Permeabilisation buffer with 5% Normal Donkey Serum at +4^0^C and 1 h staining with DyLight^TM^ 649 Donkey anti-rabbit IgG (BioLegend, #406406) diluted 1:500 in 1x Permeabilisation buffer, and fluorescence intensities were measured on Cytek® Aurora as above. The unmixing and compensation were performed in SpectraFlo®, and the data were further analysed using FlowJo v10.8.1.

For LRRK2 activity measurements in human peripheral blood neutrophils and PBMCs, the isolated cells were transferred into 96-well plate and aliquoted at 2 × 10^6^ density per condition. Cells were resuspended in 200µl of PBS and washed twice by centrifugation at 1500 rpm for 3 min at room temperature (RT). Subsequently, the cells were stained with viability dye, Live/Dead fixable stain (Thermo Fisher Scientific, # L23105) diluted in PBS (1:1000) and incubated in the dark for 20 min at RT. Following the incubation, the cells were washed twice with PBS as described above and the cell pellet was resuspended into 25 µl of FACS buffer (1% FBS in PBS) containing human FC block (BD Bioscience, # 564219; 1:200 dilution) and either 200 nM MLi-2 (synthesised by Natalia Shpiro, University of Dundee) or an equal amount of vehicle, 0.1% DMSO. Samples were incubated in the dark for 30 min at RT. To identify CD14+ monocyte, naïve B cell, and T cell populations in PBMCs, cells were stained with a cocktail of fluorophore conjugated cell surface markers, including CD3 (Biolegend, Cat# 300308), CD19 (Biolegend, # 302230) and CD14 (Biolegend, # 367112) diluted in FACS buffer (1:50 dilution) for further 30 min in the dark at RT. Following the incubation, the cells were washed twice with PBS (200 µl) and centrifuged at 1500 rpm for 3 min. Cells were fixed in 100 µl of 2% paraformaldehyde (PFA, Thermo Fisher Scientific, #J19943.K2) diluted in PBS at RT in the dark for 20 min. Subsequsently, 100 µl of working permeabilisation buffer (diluted 1:10 in MQ water, Thermo Fisher scientific, # 00-8333-56) was added to the samples followed by 3 min centrifugation at 2000 rpm at RT. Cells were washed twice with 200 µl of working permeabilization buffer as described above. Anti-pT73 Rab10 antibody (clone MJF-R21-22-5, AbCam ab241060) was diluted in working permeabilisation buffer containing 5% Normal Goat Serum (NGS, Abcam, Cat# ab7481) at 1:500 concentration and cells were incubated at 4°C overnight. Following incubations, samples were washed three times in working permeabilization buffer (by centrifugation at 2000rpm, 3 min, RT) and stained with 50 µl of Alexa Fluor 647 (Thermo Fisher Scientific, Cat# A-21245) goat anti-rabbit secondary antibody (1:500, diluted in working permeabilisation buffer containing 5% NGS). Samples were incubated for 1 hour at RT protected from light. Cells were washed with working permeabilisation buffer as described above (2000 rpm, RT, 3 min) followed by 2 washes in 200 µl of PBS. Samples were resuspended in 300µl of FACS buffer, and the data was acquired using BD LSR Fortessa with a minimum of 50,000 event acquisition per sample. The data were further analysed using FlowJo v10.8.1. LRRK2 activity in human cells was defined as:

LRRK2 activity = (pRab10 gMFI_DMSO_ – sec ab gMFI_DMSO_) - (pRab10 gMFI_MLi-2_ – sec ab gMFI_MLi-2_),

where gMFI_DMSO_ or gMFI_MLi-2_ are geometric mean fluorescence intensities of samples incubated with DMSO or MLi-2, respectively

Anti-mouse flow cytometry antibodies:

| Antibody against | clone | source | a) | b) | c) | d) | e) |
| --- | --- | --- | --- | --- | --- | --- | --- |
| CD45 | 30-F11 | BD Biosciences | + | + | + |  |  |
| MHCII (I-A/I-E) | M5/114.15.2 | BioLegend | + | + | + |  | + |
| CD11c | N418 | BioLegend | + | + | + |  |  |
| CD11b | M1/70 | BioLegend | + |  |  |  |  |
| Ly6G | 1A8 | BioLegend | + |  |  |  |  |
| B220 | RA3-6B2 | BioLegend | + |  |  |  | + |
| TCRb | H57-597 | BD Biosciences | + |  |  |  |  |
| SiglecF | E50-2440 | BD Biosciences | + | + |  |  |  |
| CD64 | S18017D | BioLegend | + | + |  |  |  |
| Ly6C | HK1.6 | Invitrogen | + |  |  |  |  |
| F4/80 | BM8 | BioLegend | + |  |  |  |  |
| NK1.1 | pk136 | BioLegend |  | + |  |  |  |
| CD3 | 17A2 | BioLegend |  | + | + |  |  |
| CD19 | 6D5 | BioLegend |  | + | + | + | + |
| EpCAM | G8.8 | eBioscience |  |  | + |  |  |
| CD93 | AA4.1 | BioLegend |  |  |  | + |  |
| IgM | RMM-1 | BioLegend |  |  |  | + |  |
| CD23 | B3B4 | BioLegend |  |  |  | + |  |
| CD21/CD35 | 7E9 | BioLegend |  |  |  | + |  |
| CD69 | H1.2F3 | Invitrogen |  |  |  |  | + |

- a) for splenocytes and lamina propria cells - b) for intraepithelial immune cells and controls (EGFP-LRRK2 levels) - c) for intraepithelial immune cells and controls (LRRK2 activity) - d) for B-cell subsets - e) for B-cell purity and activation

### Immunoblotting

Cell pellets were washed in PBS, lysed in iced-cold lysis buffer (50 mM Tris–HCl, pH 7.4, 1% Triton X-100, 10% (by vol) glycerol, 150 mM NaCl, 1 mM sodium orthovanadate, 50 mM sodium fluoride, 10 mM 2-glycerophosphate, 5 mM sodium pyrophosphate, 1 mg/ml microcystin-LR, and cOmplete EDTA-free protease inhibitor cocktail (Roche, #11836170001) and clarified by centrifugation at 17000g for 15 min at + 4°C. Frozen mouse tissues were transferred into Precellys tubes and lysed/homogenised in the above lysis buffer in the pre-chilled Cryolys Evolution at 6800 rpm, with 3 x 20 sec cycles and 2 x 30 sec pauses, and clarified by centrifugation at 17000 g for 20 min. Protein concentration in soluble lysate fraction was measured by Bradford assay (Bio-Rad Protein Assay Dye Reagent Concentrate, #5000006). The samples were prepared in 1x (final concentration) lithium dodecyl sulphate sample buffer (NuPage LDS Sample buffer Invitrogen, #NP0008) and heated at 95°C for 5 min. 20 µl (for MODE-K cells) or 37 µg (for splenocytes) of protein, or the amount of lysate corresponding to 3×10^5^ cells (for B-cells), was loaded per well onto NuPAGE 4–12% Bis–Tris Midi Gel (Thermo Fisher Scientific, Cat# WG1403BOX) and separated under NuPAGE MOPS SDS running buffer (Thermo Fisher Scientific, # NP0001-02). Proteins were transferred onto a nitrocellulose membrane (GE Healthcare, Amersham Protran Supported 0.45 mm NC) at 90V for 90 min on ice in transfer buffer (48mM Tris–HCl and 39 mM glycine supplemented with 20% methanol). The membrane cut into appropriate fragments was blocked for 1 h with 5% (w/v) skimmed milk powder in TBS-T (20 mM Tris–HCl, pH 7.5, 150 mM NaCl and 0.1% (v/v) Tween 20) at room temperature. The membrane was further incubated with rabbit anti-LRRK2 pS935 (UDD2, produced in house), anti-LRRK2 C-terminus total (NeuroMab, clone N241A/34, #75-253), anti-α-tubulin (Cell Signaling Technology, clone DM1A, mAb #3873), anti-total Rab10 (either Abcam, #ab237703, or nanoTools, clone 605B11, #0680/10), anti-phospho-Thr73-Rab10 (AbCam, #ab230261), anti-phospho-Tyr641-STAT6 (Cell Signaling Technology, #9361), anti-Beta Actin (Proteintech, #66009-1-Ig) or anti-glyceraldehyde-3-phosphate dehydrogenase (GAPDH, Santa Cruz Biotechnology, #sc-32233) antibody overnight at 4°C. Most of primary antibodies with known concentration were used at 1 μg/ml final concentration, except for total Rab10 (nanoTools) which was used at a concentration of 2 µg/ml, or diluted 1:1000, and incubated in TBS-T containing 5% BSA with exception of α-tubulin, β-actin and GAPDH antibodies that were diluted 1:5000, 1:5000 and 1:2000, respectively. After three washes with TBS-T for 10 min each, membranes were incubated with respective species-appropriate goat anti-mouse IRDye 680LT (LI-COR, #926-68020), goat anti-mouse IRDye 800CW (#926-32210) or donkey anti-rabbit IRDye 800CW (LI-COR, #926-32213) secondary antibody diluted in TBS-T (1:10000 dilution) for 1 h at room temperature, in the dark. Membranes were washed with TBS-T three times with a 10 min incubation for each wash and kept in the dark. Protein bands were acquired via near infrared fluorescent detection using the Odyssey CLx imaging system and quantified using the Image Studio software.

### Immunoprecipitation

EGFP-Lrrk2-KI lung tissues were lysed/homogenised as above with the lysis buffer containing 20 mM HEPES, pH = 7.5, 150 mM NaCl, 0.5% NP-40 supplemented with 10 mM sodium pyrophosphate, 1 mM sodium orthovanadate, 0.5 µg/ml Microcystin-LR and 1x protease inhibitor cocktail. 4 mg of tissue extracts were incubated with 25 µl of packed GFP-trap beads (Chromotek) for 2 hours before beads were isolated from the unbound fraction on the magnet, washed three times in Wash Buffer (20 mM HEPES pH 7.5, 150 mM NaCl, 0.1% NP-40 and protease/phosphatase inhibitors), and heated in 2xLDS Sample buffer at 70^0^C for 15 min. The eluted proteins were separated from beads using top speed centrifugation through a 0.22 μm Spinex column and 10% (v/v) of 2-mecraptoethanol added to eluate.

### RT-PCR

RNA was prepared from pelleted cells or frozen ileal tissues using PureLink^TM^ RNA Mini Kit (Invitrogen by Thermo Fisher Scientific, #12183025) following manufacturer’s instructions. RNA was converted to cDNA using PrimeScript^RT^ reagent Kit with gDNA Eraser (Takara, #RR047A), and used as a template for qPCR with TB Green® Premix Ex TaqTM II (Tli TNaseJ Plus) (Takara, #RR820L) and the following primers (Eurofins):

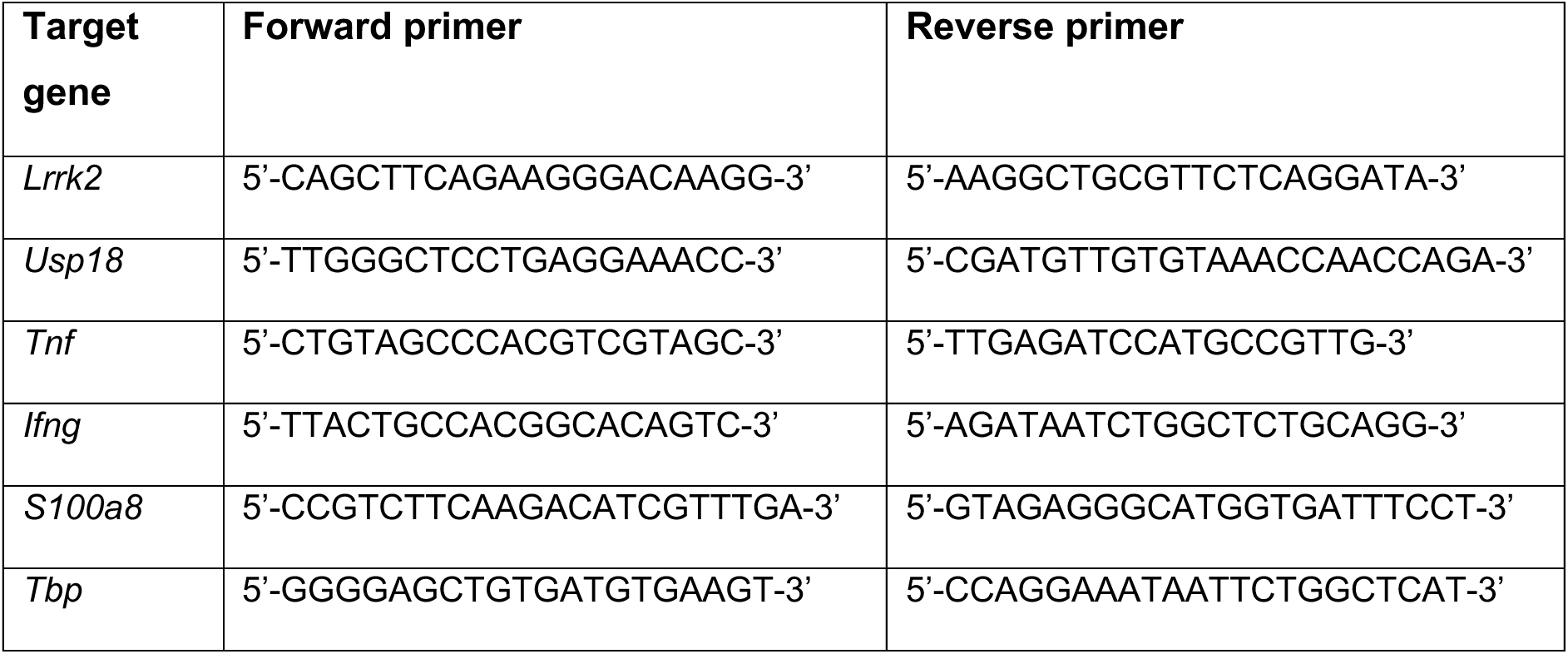

qPCR was performed in triplicates on thermocycler CFX Opus 384 (Bio-Rad Laboratories) operated by CFX Maestro software (Bio-Rad) using cycling protocol of 30 sec at 95^0^C followed by 40 cycles of 5 sec at 95^0^C and 60 sec at 60^0^C, and followed by melting curve from 65^0^C to 95^0^C. The fold change in expression was calculated using 2^-DDCT^ method (Livak & Schmittgen, 2001).

### Statistical analysis

Data were analysed in R or Microsoft Excel using one- or two-way ANOVA (with Tukey post-hoc analysis) or t-test as indicated. Analysis of Variance (ANCOVA) was performed in R.

## Acknowledgements

We are grateful for the support and helpful discussions with Dr. Hart Rardin and Dr. Yao Wong from Interline Therapeutics. We thank Dr Natalia Shapiro for producing MLi-2. We would like to acknowledge the Resource Unit at the University of Dundee for maintaining the mice and Dr. Amanpreet Singh Chawla, University of Dundee, for intraperitoneal injections, the Flow Cytometry and Cell Sorting Facility at the University of Dundee for single cell sorting, Dundee Imaging Facility at the University of Dundee for processing tissue samples and H&E staining, Tom MaCartney for LRRK2-targeting CRISPR-Cas9 plasmids and the Swamy lab for help with processing tissues. We also acknowledge Dr. Purbasha Bhattacharya for critical reading of the manuscript and Prof. Simon Arthur, University of Dundee, for helpful advice on B-cells. We also thank the volunteers who donated blood for this research.

## Author contributions

MS and DRA conceived and conceptualised the study and obtained funding. DD, RP, MT, AT, KZ and AJMH performed experiments and analysed data, DD prepared the figures and DD and MS wrote the manuscript and input from other authors. ES and AJMH provided intellectual input.

## Data availability

All data presented in this study are shown in the figures. Raw data can be made available on request.

