## SUPPLEMENTAL FIGURES for "Regulation of Leucine-Rich Repeat Kinase 2 by inflammation and IL-4"

### Regulation of Leucine-Rich Repeat Kinase 2 by inflammation and IL-4
### Supplementary data

Dina Dikovskaya[1,2]*, Rebecca Pemberton[1], Matthew Taylor[1,3], Anna Tasegian[1,4], Karolina Zenevicuite[1], Esther Sammler[1,5], Andrew J.M. Howden[6], Dario R. Alessi[1], Mahima Swamy[1]*

[1] Medical Research Council (MRC) Protein Phosphorylation and Ubiquitylation Unit, School of Life Sciences, University of Dundee, Dow Street, Dundee DD1 5EH, United Kingdom

[2] Current address: Peninsula Medical School, University of Plymouth, Drake Circus, Plymouth, PL4 8AA, United Kingdom

[3] Current address: GlaxoSmithKline, Stevenage, United Kingdom

[4] Current address: Amphista Therapeutics Ltd., Granta Park, Great Abington, Cambridge, CB21 6GQ, United Kingdom

[5] Molecular and Clinical Medicine, Ninewells Hospital and Medical School, University of Dundee, Dundee DD1 9SY, UK

[6] Cell Signalling and Immunology, School of Life Sciences, University of Dundee, Dow Street, Dundee DD1 5EH, United Kingdom

Dr. Mahima Swamy,

### Suppl Figure 1

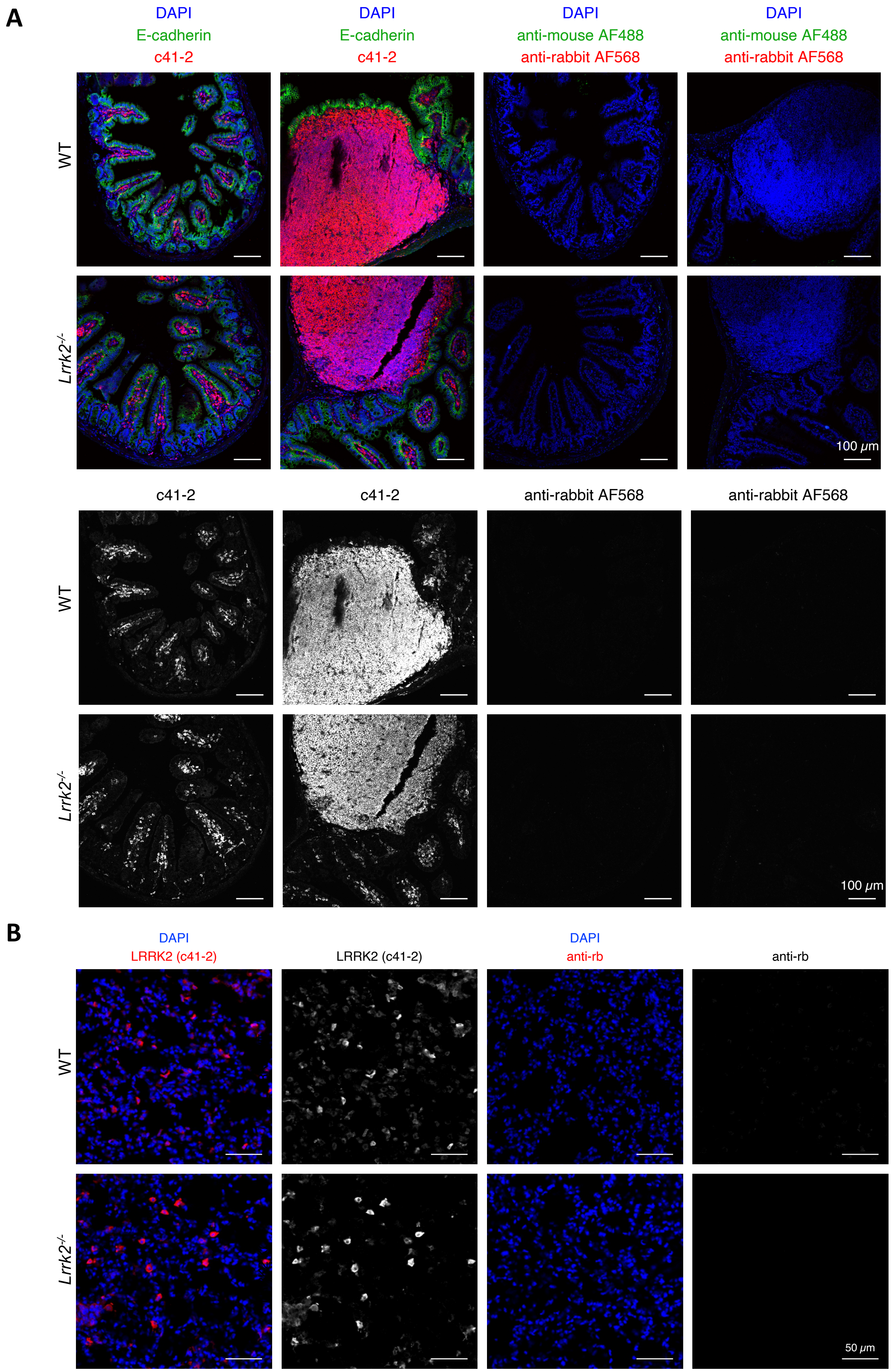

**Suppl. Figure 1.** Tissue immunofluorescence with anti-LRRK2 antibody c41-2. **A**. Sections from paraffin-embedded ileum from a C57Bl/6J (WT, rows 1 and 3) or *Lrrk2*$^{-/-}$ (rows 2 and 4) mouse were co-stained with rabbit c41-2 anti-LRRK2 ab [clone MJFF2], red, and mouse anti-E-cadherin ab (green), followed by secondary anti-rabbit Alexa Fluor-568 (AF568) and anti-mouse Alexa Fluor-488 (AF488) antibodies, counterstained with DAPI (blue) and imaged by confocal microscopy (panels 1 and 2 from left). Staining that did not include primary antibodies used as a control (panels 3 and 4) was processed, imaged and adjusted in in the same way. Single optical sections are shown. Panels 2 and 4 depict areas with Peyer's Patches. Top two rows show overlay of all staining, and the bottom two rows show only c41-2 channel from the same images in black and white. Size bars are 100 µm. **B.** Sections from frozen lung tissues from WT (top) or *Lrrk2*$^{-/-}$ (bottom) mice were stained with c41-2 anti-LRRK2 ab (clone MJFF2) (red on overlay) followed with Alexa Fluor-568 fluorescent secondary ab (panels 1 and 2), or with secondary ab only (panels 3 and 4). Sections were counterstained by DAPI (blue) and Phalloidin-488 (not shown). Confocal images were acquired and processed identically. Panels 1 and 3 show c42-1 and DAPI overlay, panels 2 and 4 show c42-1 channel only in black and white. Size bars are 50 µm. Images were processed and assembled in OMERO.

### Suppl. Figure 2

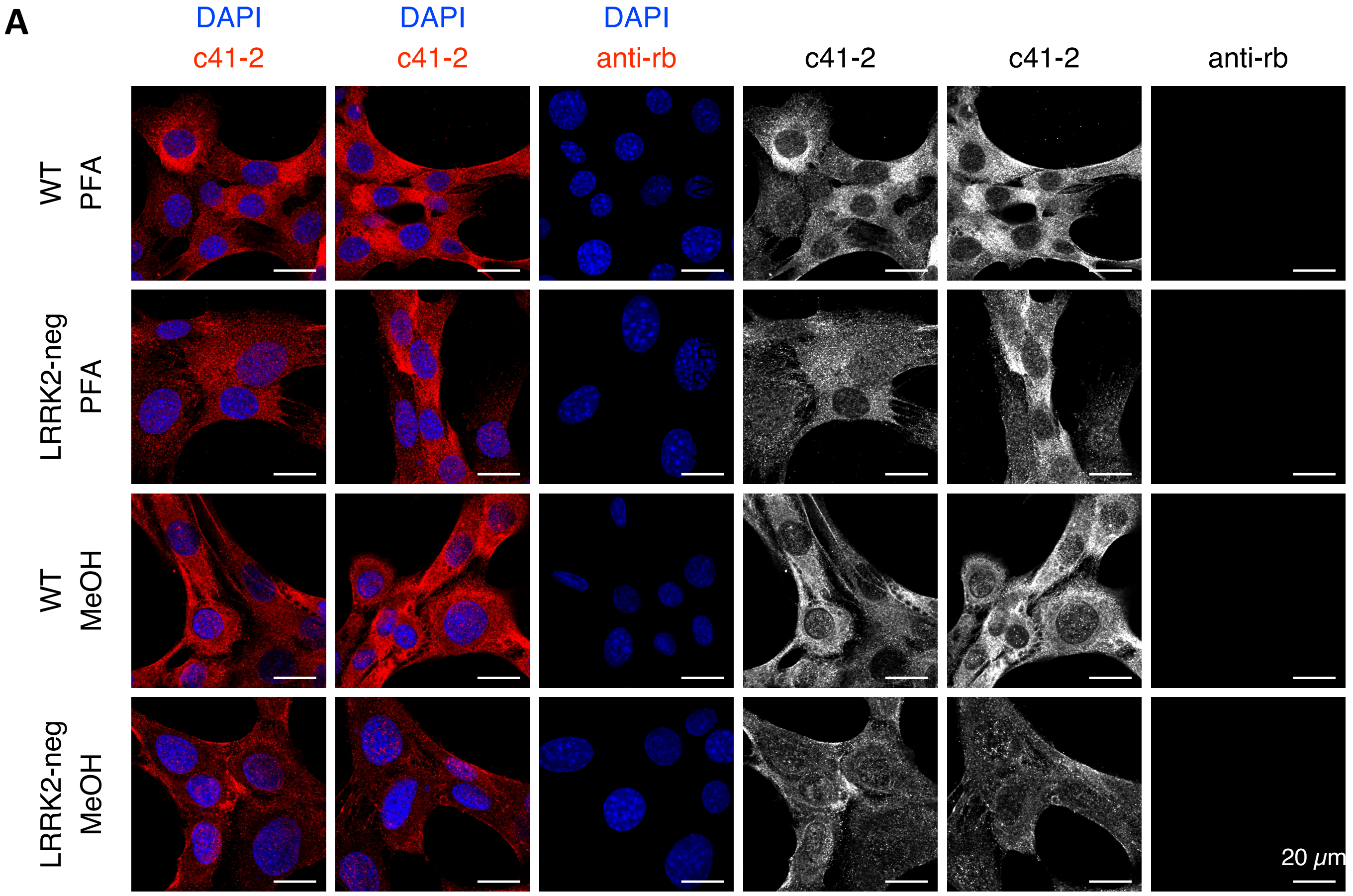


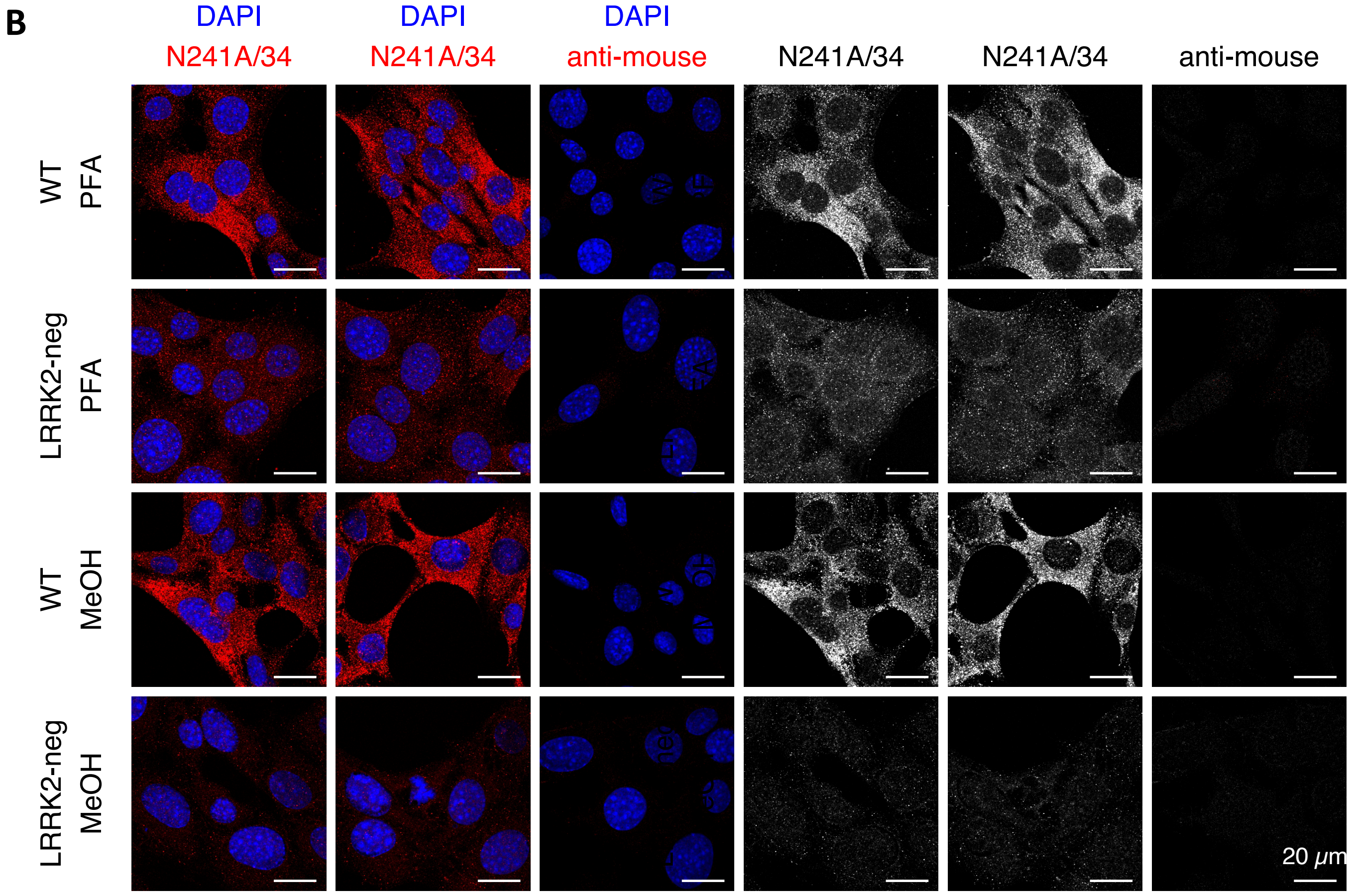

**Suppl. Figure 2.** Validation of anti-LRRK2 immunofluorescence using MODE-K cells. Parental MODE-K cells (WT) and the LRRK2-deficient clone (LRRK2-neg) generated by CRISPR-Cas9 knock-out of LRRK2 (see Fig 1A for Western blot) were grown on coverslips, fixed/permeabilised in either 4% PFA/1% TritonX100 (top two rows, PFA) or Methanol (bottom two rows, MeOH), and stained with either c41-2 (clone MJFF2), **(A)** or clone N241A/34 **(B)** anti-LRRK2 antibodies  followed by an appropriate fluorescently-labelled secondary ab (red on overlay) and DAPI (blue on overlay). Cells were imaged on confocal microscope and images processed in OMERO. Anti-rabbit (anti-rb) or anti-mouse secondary-only controls (panels 3 and 6) were imaged and processed identically to the full-stained samples. The overlays are shown in panels 1-3, and panels 4-6 depict LRRK2-only staining in black and white. Size bars 20 μm.

### Suppl. Figure 3

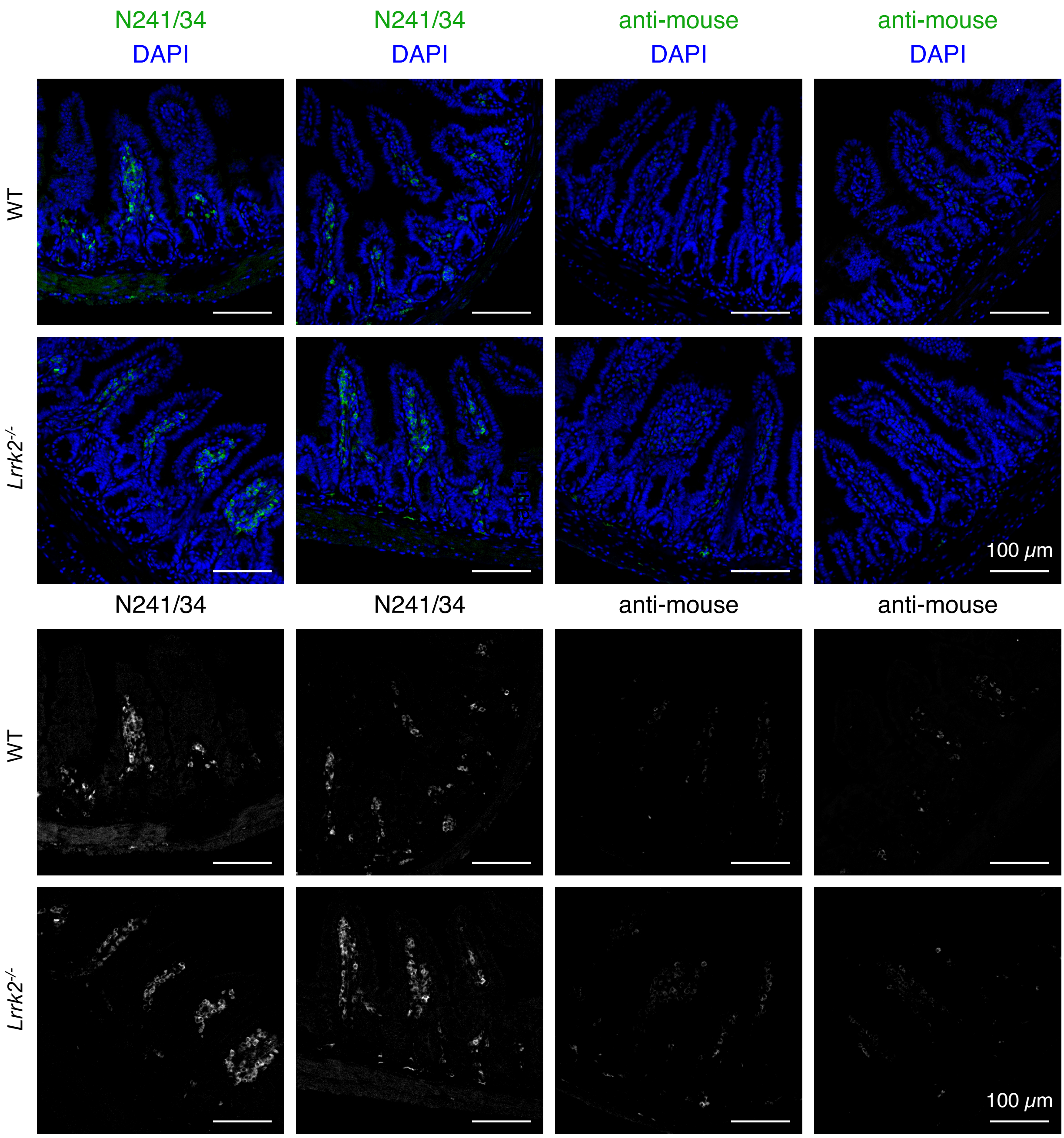


**Suppl. Figure 3.** Tissue immunofluorescence with anti-LRRK2 antibody N241A/34. Sections of paraffin embedded ileums from *Lrrk2*$^{-/-}$ mouse (rows 2 and 4) and its WT littermate (rows 1 and 3) were stained with N241A/34 antibodies followed by fluorescent secondary ab (panels 1 and 2) or with secondary ab only (panels 3 and 4), (green on overlay) and counter-stained with DAPI (blue on overlay). Tissues were imaged on confocal microscope and processed in OMERO in the same way. Top two rows show overlay, bottom two rows show LRRK2-only staining in black and white. Size bars 100 μm.

### Suppl. Figure 4

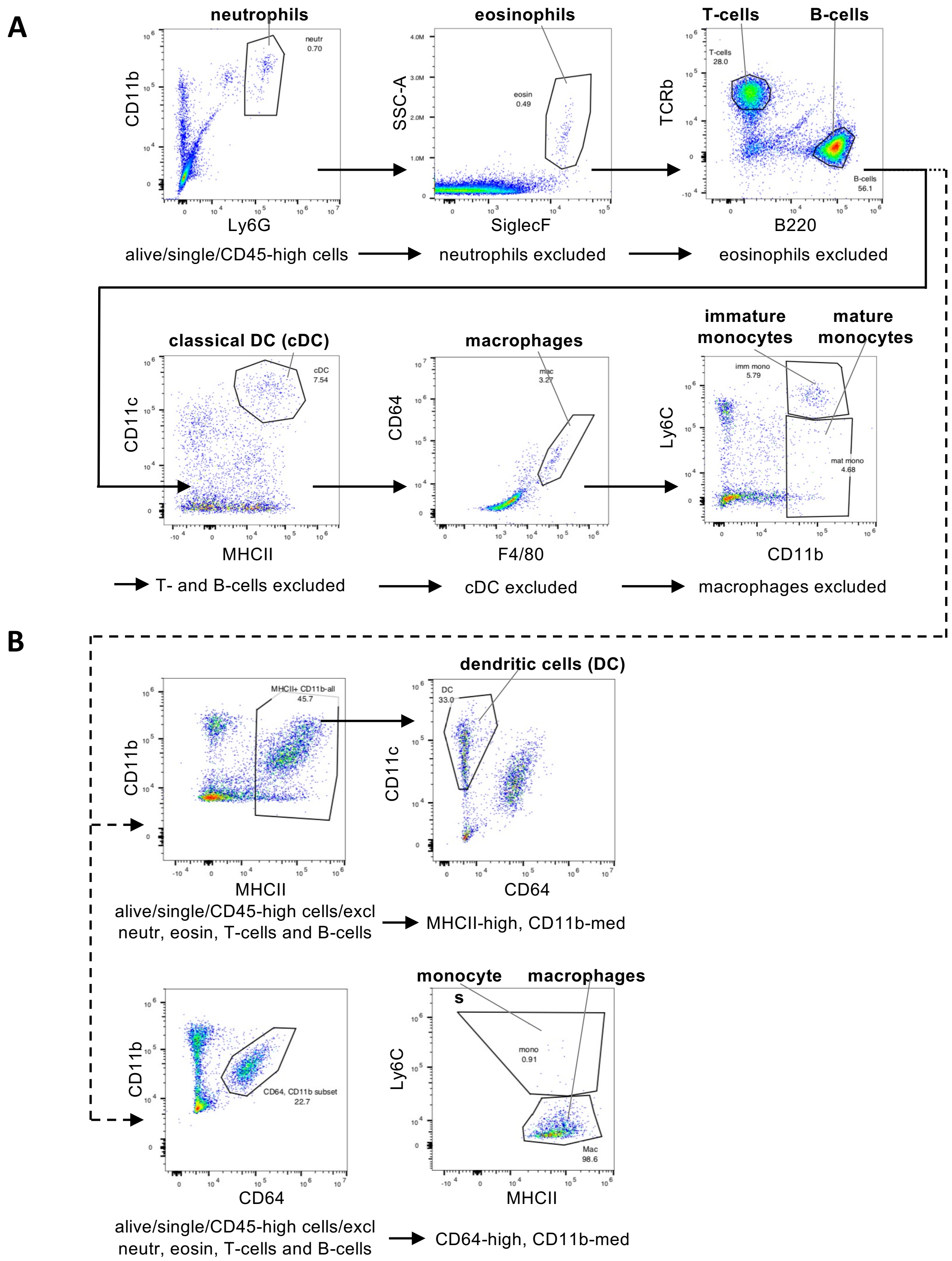


**Suppl. Figure 4.** Gating strategy for immune cell types. A. Splenocytes gating. Immune cells were gated from live, single CD45+ cells followed by sequential identification and exclusion of: neutrophils ($CD11b^{+}/Ly6G^{hi}$); eosinophils ($SiglecF^{+}$/ SSC-high); T- cells (TCRb+) and B-cells (B220-high); classical dendritic cells (cDC, $CD11b^{hi}/MHCII^{hi}$); macrophages (CD64+/F4/80+) and immature and mature monocytes ($CD11b^{hi}/Ly6C^{hi}$ and $CD11b^{hi}/Ly6C^{lo}$ respectively). **B.** Lamina propria cells gating. The staining and initial gating was identical to A up to the exclusion of T- and B-cells, after which DCs were defined as $CD11b^{+/-}/MHCII^{hi}$ then gating on $CD11c^{hi}/CD64^{-}$ cells. Macrophages and monocytes were identified as CD64+/ $CD11^{int}$ cells, then $Ly6C^{+}$ for monocytes and for $Ly6C^{-}/MHCII^{+}$ for macrophages.

### Suppl. Figure 5

**A**

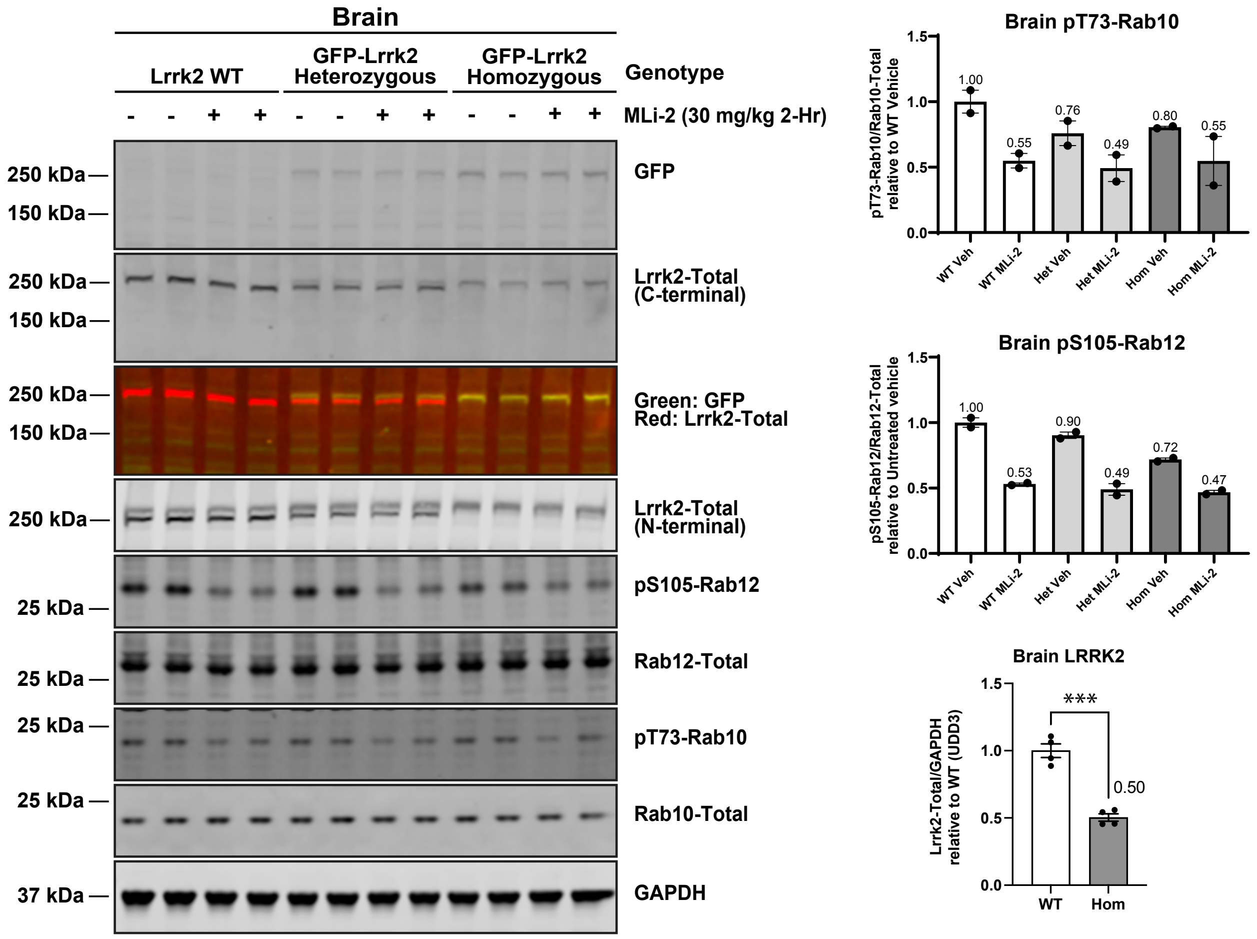


**B**

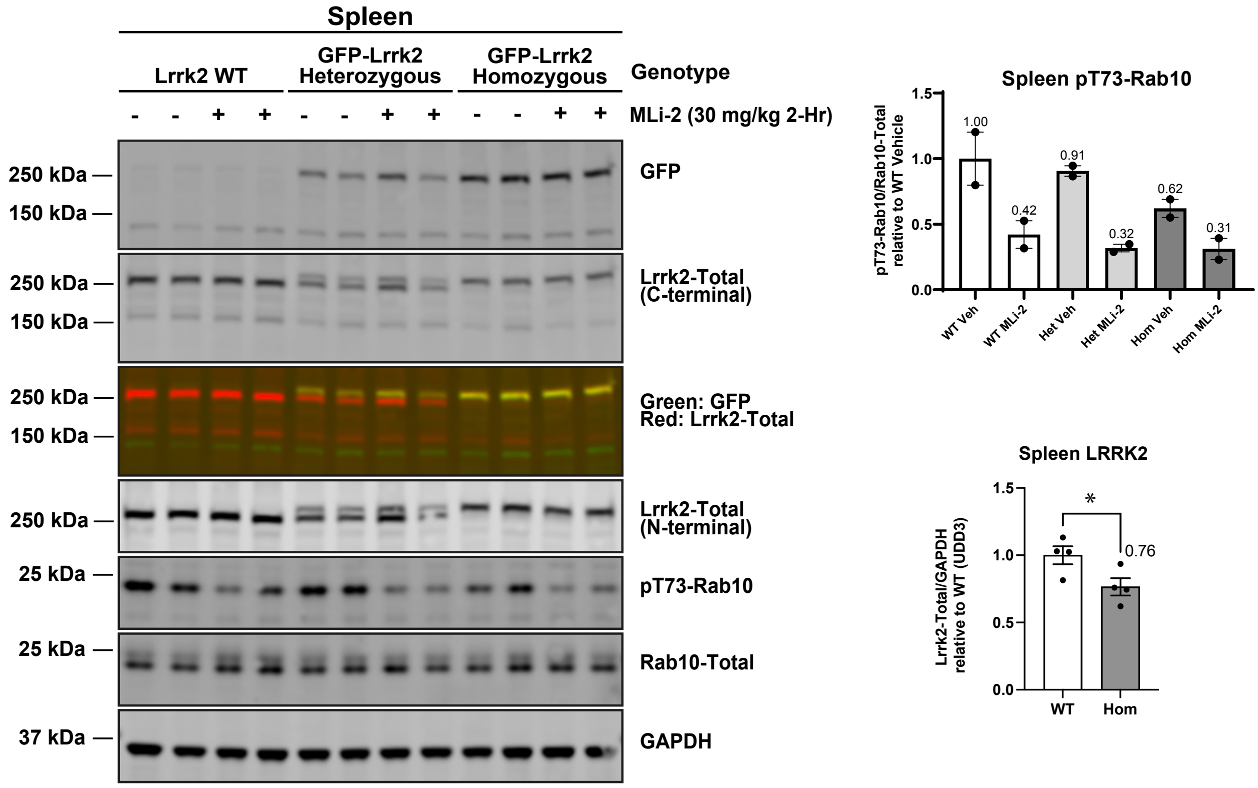

### Suppl. Figure 5

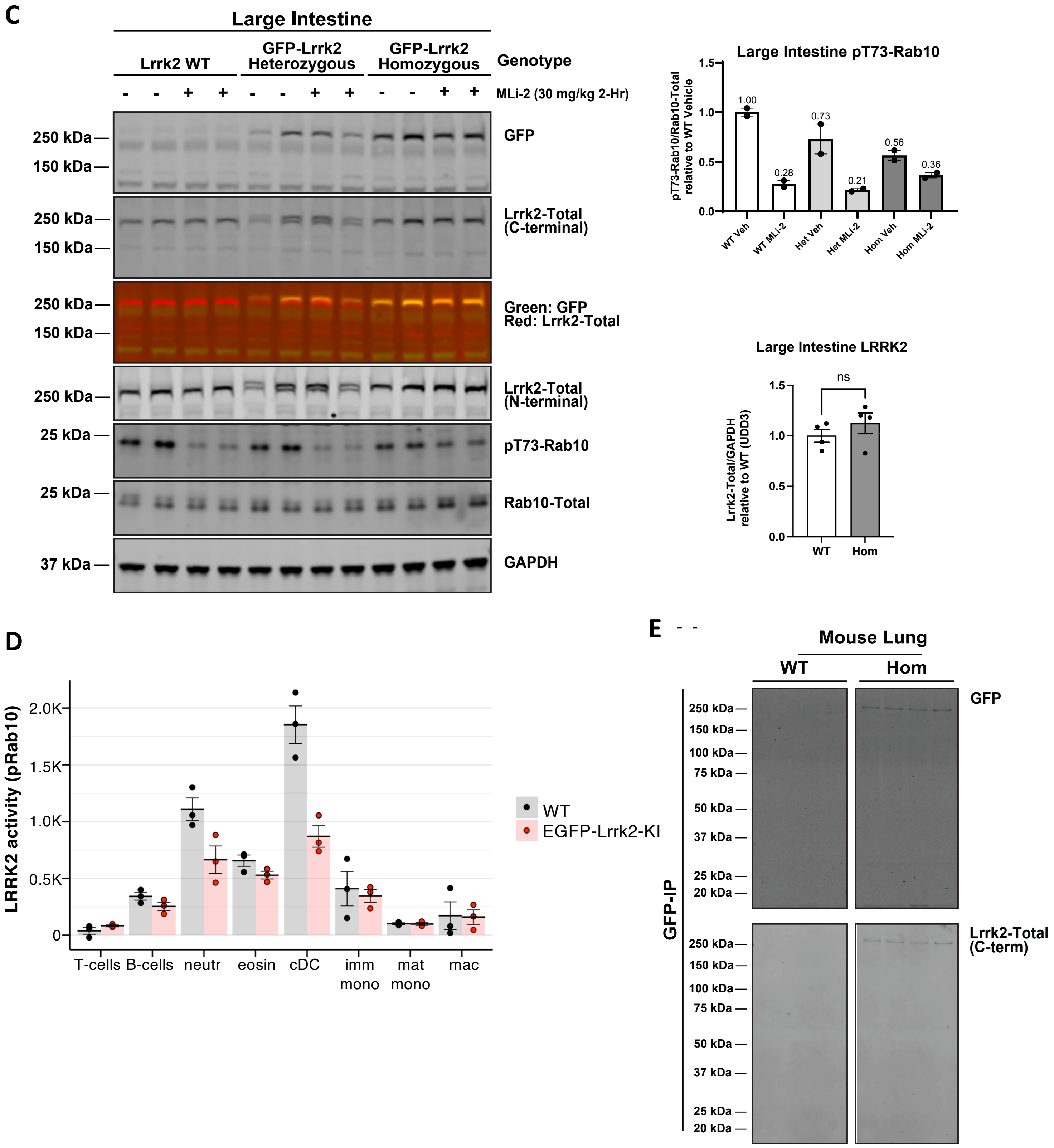


**Suppl Figure 5.** Analysis of Lrrk2 pathway in tissues from EGFP-Lrrk2-KI mouse model. A-C. 3-month-old WT, Heterozygous and Homozygous EGFP-Lrrk2-KI mice were treated with or without 30 mg/kg MLi-2 for 2h prior to culling. Mouse tissues were immediately extracted and snap frozen in liquid nitrogen. Frozen tissues were lysed using Cryolys Evolution and between 20 to 30 µg of whole tissue lysate of either brain (A), spleen (B) or large intestine (C) subjected to immunoblot analysis. (contd. on next page)

**Suppl Figure 5 (contd).** Quantification of Lrrk2-substrate phosphorylation of pT73-Rab10 and/or pS105-Rab12 relative to total levels, and total Lrrk2 levels relative to GAPDH for both N- and C-termini antibodies are shown as mean ± SEM for each tissue. Each lane indicated sample derived from a different mouse tissue. **D**. LRRK2 activity was measured and displayed as in Figure 1C in splenocytes from homozygous EGFP-Lrrk2-KI mice (EGFP-Lrrk2-KI, red) or their wild type littermates (WT, black). **E**. Mouse lung tissues derived from homozygous genotypes of Lrrk2-WT and EGFP-Lrrk2 mice were homogenised using Cryolys Evolution in 0.5% NP-40 detergent lysis buffer. 4mg of lung tissue lysate was subjected to a GFP IP at 4°C for 2h. 10% of the IP elute was subjected to immunoblot analysis using antibodies indicated.

### Suppl. Figure 6

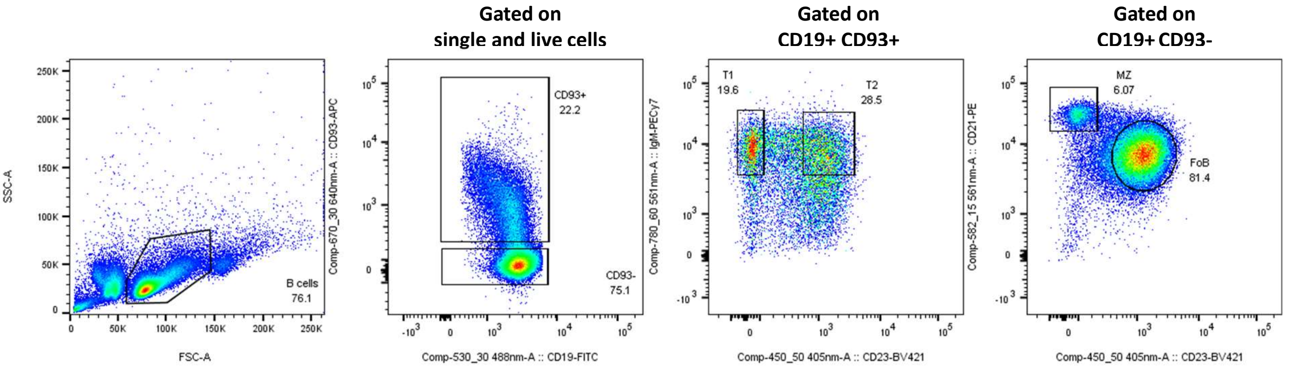


**Suppl. Figure 6.** Gating strategy for mouse splenic B-cell subtypes. Splenic B-cells were isolated using pan B-cell isolation kit by negative selection and stained with DAPI and antibodies for surface markers. Live single $CD19^{+}$ B-cells were digitally separated into $CD93^{hi}/CD23^{-}$ T1 transitional B-cells and $CD93^{hi}/CD23^{hi}$ T2 transitional B-cells, and $CD93^{-}$ mature B-cells were further subdivided $CD23^{hi}/CD21^{+}$ follicular B-cells (Fo) and $CD23^{-}/CD21^{hi}$ marginal zone B-cells (MZ).

#### Suppl. Figure 7

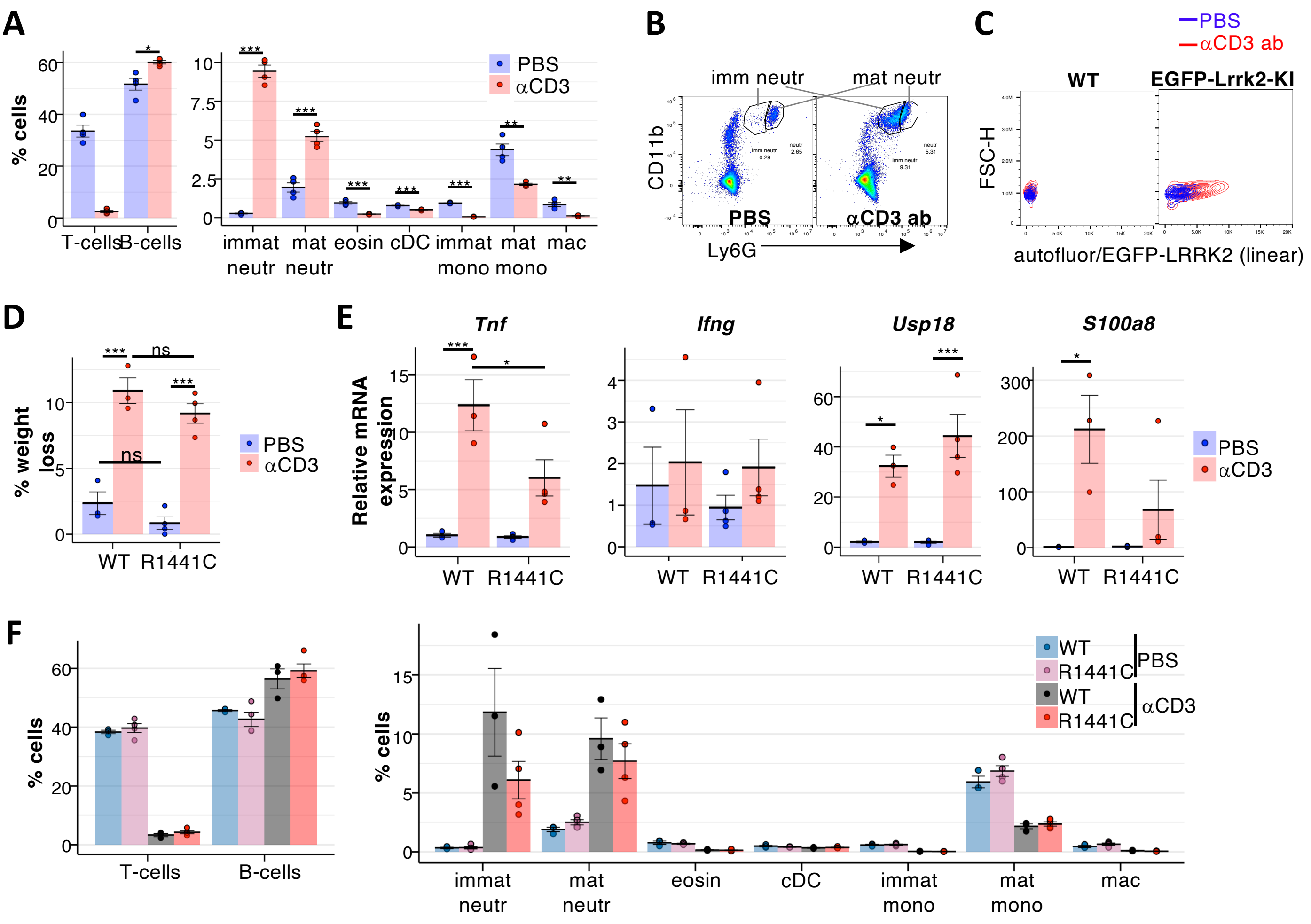


**Suppl. Figure 7.** Effects of intraperitoneal administration of anti-CD3 antibody in wild type and *Lrrk2-R1441C* mutant mice. **A.** Cellular composition of mouse spleen 24h after ip injection of anti-CD3 ab (αCD3, red) or PBS (blue). Dots show frequencies of indicated cell types in individual mice (4 mice per group) as a percent of single live $CD45^{+}$ cells. Bars show means, and error bars represent SEM. Statistical significance of the changes calculated by one-way ANOVA is shown as stars: *** ($p<0.001$) or ** ($0.001<p<0.01$). Statistical analysis was not applied to T-cells, since strong reduction in the number of T-cells is most likely due to incomplete identification of T-cells resulting from anti-CD3 ab induced internalisation of the TCR. **B.** Distinction between mature and immature neutrophils by Ly6G level from cells in A. Note a strong expansion of the $CD11b^{hi}/Ly6G^{int}$ immature neutrophil subset in anti-CD3 ab injected sample. **C.** EGFP-LRRK2 fluorescence (for EGFP-Lrrk2-KI sample) or autofluorescence measured in the same channel (for matching WT control) in B-cells among splenocytes from WT (left panel) or EGFP-Lrrk2-KI (right panel) mice 24h after anti-CD3 ab (red) or PBS (blue) injection plotted on a linear scale against FSC-H. **D.** Body weights before and 24h after ip injection of anti-CD3 antibody (aCD3 ab, red) or equal volume of PBS (PBS, blue) were measured, and % weight loss was calculated and displayed as dots (individual mouse measurements), bars (means) and error bars (SEM). Statistical significance was calculated by two-way ANOVA. **E.** Relative expression of *Tnf*, *Ifng*, *Usp18* and *S100a8* mRNA was measured by qPCR in ileums isolated from *Lrrk2-R1441C* mice (R1441C) and their wild-type littermates (WT) 24h after anti-CD3 ab (red) or PBS (blue) injection. Graphs show fold changes relative to PBS-injected WT mice, with *Tbp* as a reference gene. Dots correspond to the values in individual mice, the bars show geometric means ± SEM. Statistically significant differences are indicated by p-values calculated by two-way ANOVA. (contd. next page)

**Suppl. Figure 7 contd.** F. Frequencies of indicated cell types among live single CD45+ splenocytes obtained from WT or R1441C mice 24h after anti-CD3 (grey or red) or PBS (purple or blue) injection are displayed as individual measurements (dots), means (bars) and SEM (error bars). Note that there was no statistically significant difference (quantified by two-way ANOVA) between two genotypes.

#### Suppl. Figure 8

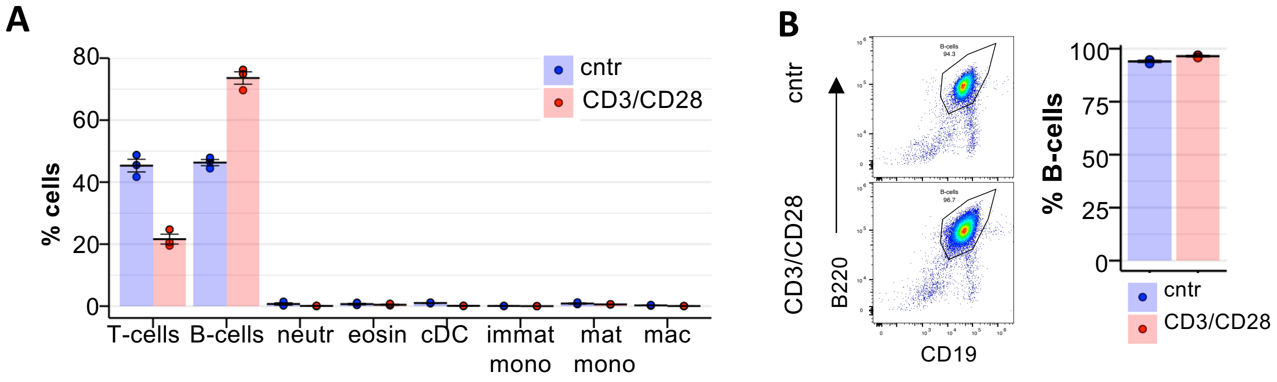


**Suppl. Figure 8.** *In vitro* B-cell stimulation by T-cell-dependent factors. **A.** Frequencies of indicated cell types detected by spectral flow cytometry (see Suppl. Fig 4A for gating) in the mixed splenocyte culture 24 h after *in vitro* incubation with immobilised anti-CD3/CD28 antibodies (CD3/CD28, red) or in control media supplemented with IL-7 (cntr, blue) were quantified as % from alive single CD45+ cells within harvested cell suspensions. **B.** Purity of B-cells isolated from mixed splenocyte cultures after 24 h incubation with immobilised anti-CD3/CD28 antibodies (CD3/CD28, bottom left panel, red on the plot) or in control media supplemented with IL-7 (cntr, top left panel, blue on plot). B-cells were purified using negative selection kit and defined as CD19+/B220+ cells (left panels). The quantification of the purity as % from single live cells is shown on the right. (A-B) Individual samples (dots), means (bars) and SEM (error bars) are shown.
